# Rapid clinical metagenomics enables early tailored therapy in complicated urinary tract infections and strengthens antimicrobial stewardship

**DOI:** 10.64898/2026.03.09.709250

**Authors:** Anurag Basavaraj Bellankimath, Isabell Kegel, Sverre Branders, Truls E. Bjerklund Johansen, Can Imirzalioglu, Torsten Hain, Florian Wagenlehner, Rafi Ahmad

## Abstract

Rapid and accurate diagnosis of UTIs remains difficult because culture-based methods are slow and less sensitive. This study evaluates URINN, a metagenomic workflow that detects uropathogens, antibiotic resistance genes, and virulence factors directly from patient urine samples. The optimized protocol was tested on a combined set of 349 clinical urine samples. URINN demonstrated 99% accuracy across all samples and 97% sensitivity for identifying 294 pathogens, including both bacteria and fungi. It predicted antibiotic susceptibility with 91% accuracy across 2099 antibiotics. The method detected pathogens at concentrations as low as 9.3 × 10^3^ CFU/mL and provided results within approximately four hours. Flow cytometry and DNA yield analyses helped establish thresholds to differentiate culture-positive from culture-negative samples, with genome coverage linked to the accuracy of susceptibility predictions for certain species. Virulence profiling revealed that adherence and nutritional factors are crucial for colonization and persistence. Leukocyte counts were comparable between genders, but bacterial loads were higher in females. The catheterized group had significantly higher leukocyte counts, and their urine showed increased cephalosporin resistance. This approach could enhance clinical decision-making, support personalized treatment, and improve the management of complicated UTIs, thereby contributing to better UTI care and antibiotic stewardship.

## Introduction

Urinary tract infections (UTIs) are among the most common bacterial infections in humans worldwide^1^. It disproportionately affects women, with 50-60% of adults experiencing it at least once in their lifetime^2^. UTIs are the main reason antibiotics are prescribed and used in adults following respiratory infections. Patients with complicated UTIs (cUTI) often experience recurrences, mainly due to risk factors such as gender, anatomical and functional urinary tract issues, urinary stones, catheter use, antibiotic exposure, and immunocompromised status state^3^. Among cUTI patients, catheter use is the main risk factor for developing catheter-associated UTI (CA-UTI)^4^. These infections are also associated with various drug-resistant bacteria that often lead to persistent and recurrent infections. This is due to their ability to form biofilms on catheters, thereby protecting them from antibiotics^5^. Currently, antibiotic treatment for UTI is usually empiric, guided by guidelines and regional resistance profiles^6^. Consequently, patients often receive multiple empiric antibiotic doses, which raises their risk of developing antibiotic resistance, leading to limited treatment options and increased mortality^7,8^. Hence, reducing overall antibiotic consumption in UTI treatment will have a significant impact on antimicrobial resistance.

Culture-based diagnostic methods for UTI have been used clinically for more than 60 years since their introduction in the early 1960s^9^. They have limitations, such as low sensitivity and long turnaround times, that impede accurate management of UTIs, particularly amid rising antimicrobial resistance. Clinical metagenomics has the potential to transform infection management by rapidly identifying all pathogens in a sample^10^. In addition to identifying pathogens and assessing resistance, clinical metagenomics offers additional data at the pathogen level. This data can be utilized for profiling virulence and abundance, conducting functional metabolic analysis, and supporting microbiome research^11,12^.

In a prior proof-of-concept study (Phase I)^13^, we demonstrated that rapid pathogen identification and antibiotic susceptibility prediction directly from urine samples using metagenomics are feasible, thereby eliminating the need for urine culture. We tested 11 methods—eight in-house and three commercial kits—on 78 UTI samples. The top performer, identified as the “optimized method,” stood out in host depletion, DNA extraction, turnaround time (TAT), and cost effectiveness. It reached 99% accuracy in pathogen identification, with 100% precision, 99% recall, and predicted antibiotic susceptibility with 90% accuracy and 95% sensitivity.

However, the study was limited by a small sample size of 78 participants, with only 53 tested using the optimized method. To address this limitation, we aimed to conduct a larger prospective study to better evaluate the robustness and diagnostic accuracy of the optimized method across a broader patient population. Thus, the main goal of this study is to refine and validate the robustness of our optimized method (referred to in this manuscript as URINN) on an additional 296 patient samples (Phase II) from patients with complicated UTI treated at a tertiary German reference center. Furthermore, we analyzed metagenomic data from the entire cohort of 349 patient samples (53 from Phase I and 296 from Phase II) to identify the most common UTI pathogens, antimicrobial resistance factors, virulence factors, and patterns of pathogen and antibiotic co-occurrence. Additionally, the data were stratified based on two risk factors, catheterization status and gender-specific risks, to allow for more detailed analysis.

## Results

### Routine microbiological results for the Phase II cohort

In the phase II cohort, a total of 296 excess patient urine samples were collected for evaluation (Supplementary Table 1). Of these, 68% (202/296) were from male patients, and 32% (93/296) from female patients, with one sample being undetermined. Among the samples, 50% (149/296) were culture-positive, of which 58% (87/149) were monomicrobial, and 42% (62/149) were polymicrobial (Supplementary Figure 1). Additionally, 142 samples were culture-negative. Overall, the most common uropathogens among the clinical samples were *Escherichia coli* (22%)*, Enterococcus faecalis* (19%), and *Klebsiella pneumoniae* (8%) (Supplementary Table 3). Of 232 bacterial pathogens, routine antimicrobial susceptibility testing (AST) was performed on 163 (Supplementary Table 4), of which 78% (132/163) were resistant to at least one antibiotic. Of the total resistant pathogens, 41% (67/143) were multidrug-resistant (resistant to three or more antibiotics).

### Minor modifications to the optimized method

To further enhance the DNA extraction protocol, minor adjustments were implemented relative to the original optimized method. During host depletion, the saponin concentration was increased from 2.2% to 3%. A SYBR-green-based qPCR assay showed no significant differences in host depletion between these two saponin concentrations and showed no loss of bacterial DNA (Supplementary Figure 2). Culture-positive samples with low biomass, totaling less than 200 ng after AmpureXP purification, were amplified using a whole-genome amplification (WGA) kit. In total, 57 samples were subjected to amplification: 44 were successfully amplified, 12 failed, and 1 was excluded due to sampling error.

### Comparison between URINN results and standard urine culture (MALDI-TOF/VITEK-2)

The URINN results showed strong agreement with routine clinical microbiology results for both pathogen identification (MALDI-TOF) and AMR detection (VITEK-2) (Table 1). It demonstrated an overall accuracy score of 99% across 296 samples (Figure 1). At the pathogen level (Table 1), the method demonstrated an overall accuracy (prevalence threshold) of 97% (257/264), with a sensitivity of 97% (199/206), specificity of 79% (58/73), negative predictive value (NPV) of 89% (58/65), and positive predictive value (PPV) of 93% (199/214). The method had six false negatives, missing the pathogens *Streptococcus mitis* (in samples 133 and 240), *Morganella morganii* (196), *Acinetobacter baumannii* (198), *Proteus mirabilis* (233), and *E. faecalis* (253). Four of these samples (196, 198, 240, and 253) were polymicrobial samples (Supplementary Figure 1). The method also identified 15 additional pathogenic bacteria in 15 tested samples (Supplementary Figure 3) that routine microbiology had failed to detect. The method also exhibited 92% accuracy (1330/1446) in predicting antibiotic susceptibility, with a specificity of 99% (1153/1169), sensitivity of 64% (177/277), precision of 92% (177/193), and NPV of 92% (1153/1253). Most false-negative/positive predictions were for the ampicillin, amoxicillin-clavulanic acid, ciprofloxacin, and trimethoprim/trimethoprim sulfamethoxazole antibiotic groups (Supplementary Figure 4 and Supplementary Figure 5).

**Figure 1:**
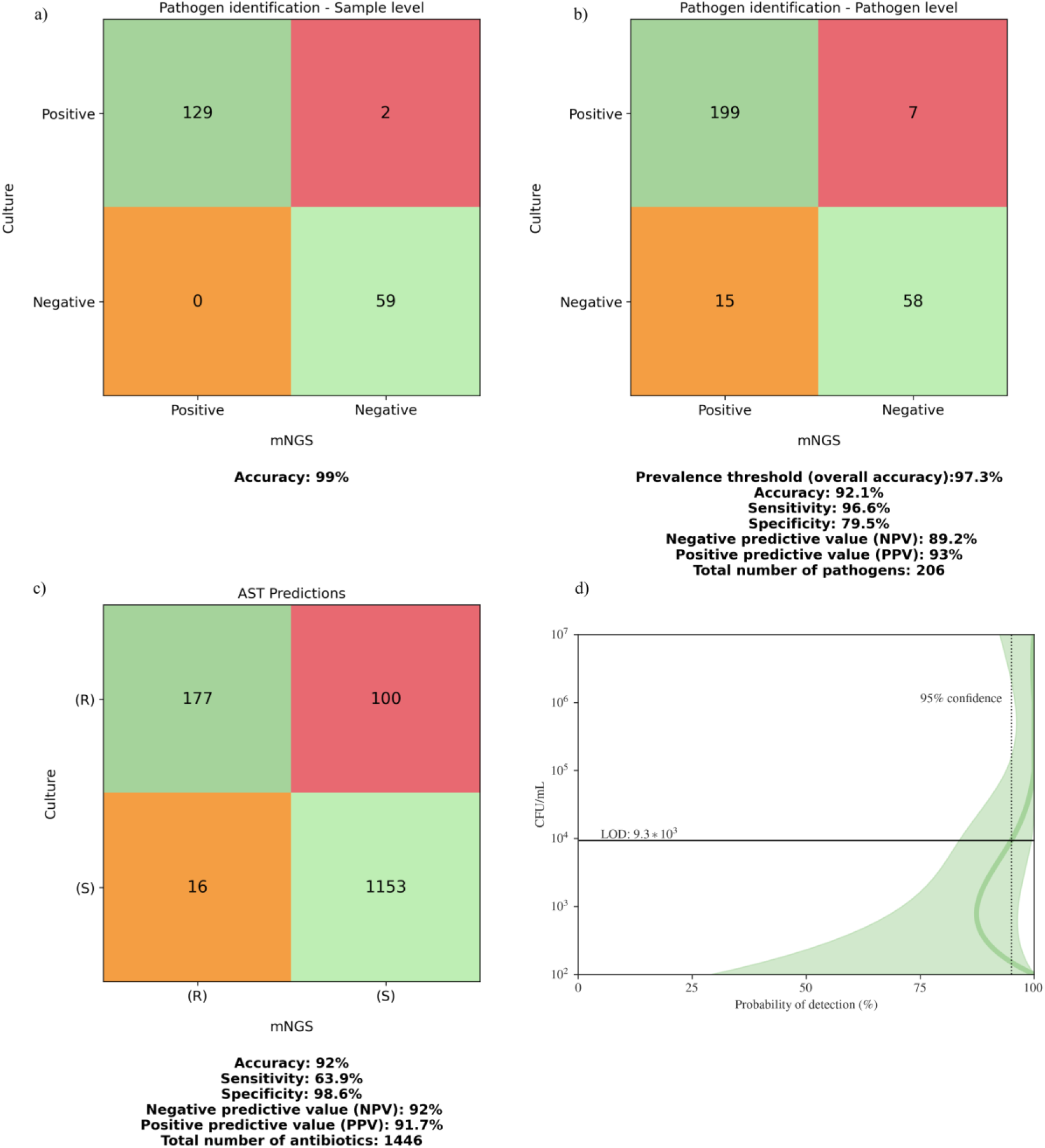
Confusion matrix representing the performance of the method for pathogen identification and antibiotic susceptibility predictions and limit of detection based on Phase II cohort: Subfigures (a) & (b) represent the pathogen identification performance of the method stratified on the sample and pathogen level, while subfigure (c) represents the antibiotic susceptibility predictions on the antibiotic level. The probability of detection at each concentration of colony-forming units (CFU) was determined using the beta-distribution approximation (d). The observed probability of detection is plotted as a function of bacterial CFU/mL. The 95% confidence interval for the probability of detection is indicated by the shaded area. The dotted line indicates the threshold for a 95% probability of detection, corresponding to a limit of detection of approximately 9.3 × 10^3^ CFU/mL.

**Table 1:** An overview of the scoring matrices obtained by comparing mNGS results with routine MALDI/VITEK-2 testing, along with a comparison of accuracy metrics between Phase I and Phase II results. Metrics for agreement between metagenomics results and routine pathogen identification and antibiotic susceptibility testing (AST) are shown for both study phases and the combined scores on a pathogen basis. Each metric is expressed as a percentage and accompanied by the absolute numbers in parentheses. The 95% confidence interval (CI) is provided for each metric as a percentage.

|  | Phase | Prevalence<br>threshold | Sensitivity<br>(TPR) | Specificity<br>(TNR) | Precision<br>(PPV) | Negative<br>Predictive<br>Value (NPV) | Accuracy<br>Score |
| --- | --- | --- | --- | --- | --- | --- | --- |
| **Pathogen**<br>**ID** | **I** | 99%<br>(100/101) CI<br>97-100% | 99%<br>(87/88)<br>CI 97-100% | 100%<br>(13/13)<br>CI 100-100% | 100%<br>(87/87)<br>CI 100-100% | 93%<br>(13/14)<br>CI 79-100% | 99%<br>(100/101)<br>CI 97-100% |
|  | **II** | 97%<br>(257/264)<br>CI 95-99% | 97%<br>(199/206)<br>CI 94-99% | 79%<br>(58/73)<br>CI 70-89% | 93%<br>(199/214)<br>CI 90-96% | 89%<br>(58/65)<br>CI 82-97% | 92%<br>(257/279)<br>CI 89-95% |
|  | **I + II** | 98%<br>(357/365) CI<br>96-99% | 97%<br>(286/294) CI<br>95-99% | 83%<br>(71/86)<br>CI 75-91% | 95%<br>(286/301)<br>CI 93-97% | 90%<br>(71/79)<br>CI 83-97% | 94%<br>(357/380) CI<br>92-96% |
| **AST** | **I** |  | 68%<br>(82/121) CI<br>59-76% | 95%<br>(507/532) CI<br>94-97% | 77%<br>(82/107) CI<br>69-85% | 93%<br>(507/546) CI<br>91-95% | 90%<br>(589/653) CI<br>88-92% |
|  | **II** |  | 64%<br>(177/277) CI<br>58-70% | 99%<br>(1153/1169)<br>CI 98-99% | 92%<br>(177/193)<br>CI 88-96% | 92%<br>(1153/1253)<br>CI 91-94% | 92%<br>(1330/1446)<br>CI 91-93% |
|  | **I + II** |  | 65%<br>(259/398) CI<br>60-70% | 98%<br>(1660/1701)<br>CI 97-98% | 86%<br>(259/300)<br>CI 82-90% | 92%<br>(1660/1799)<br>CI 91-94% | 91%<br>(1919/2099)<br>CI 90-93% |

For the combined cohort of 349 UTI samples from phase I and phase II, overall pathogen identification accuracy (prevalence threshold) was 98% (357/365), with 97% (286/294) sensitivity, 83% (71/86) specificity, 95% (286/301) PPV, and 90% (71/79) NPV. Antibiotic susceptibility prediction accuracy was 91% (1919/2099) overall, with 98% (1660/1701) specificity.

### MysteryMaster recovered approximately 21% of unclassified reads

Overall, 21% (2.2M/10.5M) of total sequencing reads remained unclassified. MysteryMaster recovered 26% (0.6M/2.2M) of all unclassified reads. On average, each barcode contained >50k reads with a median N50 of 4470 bp.

### Turnaround time (TAT) is approximately the same as previously reported

Minor modifications to the lysis step of the protocol did not significantly impact the TAT for the samples. Consistent with the proof-of-concept study, most samples exhibited a TAT of approximately 4 hours from the start of sample processing. The only exception was the 38 samples that required additional amplification, which extended the TAT by 90 minutes, bringing the total to 5.5 hours.

### Routine microbiological findings emphasize the complexity of the phase I and phase II cohorts

Because the current work builds on our findings from a previous proof-of-concept study at the same tertiary care center, combined data from both studies were included in the detailed analysis presented in the results that follow (flow cytometry and DNA yield AUROC analysis, limit of detection (LOD), virulence factors (species-specific), and co-occurrence analysis). The overall dataset includes 349 patient samples (Supplementary Table 1 and Supplementary Figure 1). The cohort comprised 56% positive and 44% culture-negative samples. Among the culture-positive samples, 53% were monomicrobial, and 47% were polymicrobial (Supplementary Figure 1).

### The URINN method can detect as low as 103 CFU/mL

A beta-distribution approximation was used to determine the limit of detection, defined as the probability of detecting *any* pathogen at a specific concentration. The beta-distribution approximation (Figure 1d) showed a limit of detection (95% probability of detection) for the URINN metagenomic method of 9.3 × 10^3^ CFU/mL. The samples ranged from 10^2^ – 10^7^ CFU/mL (Supplementary Figure 6). Additionally, in sample 97, the routine detected *Enterococci* species at 10^2^ CFU/mL, which the current method accurately identified as *E. faecalis*. Of the seven pathogens not detected by the URINN method (Supplementary Figure 3, Supplementary Figure 6), four had a CFU/mL of 10^3^, two had 10^4^ CFU/mL, and one had 10^5^ CFU/mL.

### The results confirm flow cytometry and DNA yield-based cutoffs for sample positivity

Flow cytometry data on bacterial and human cell counts (including erythrocytes, leukocytes, and round and squamous epithelial cells) from clinical samples (Supplementary Table 5) were used to estimate host depletion by correlating cell counts with bacterial and human read counts (Supplementary Figure 8). The URINN method accurately identified pathogens at a minimum bacteria-to-host cell ratio of 0.1. An AUROC analysis was performed to evaluate whether total DNA yield and/or the number of bacterial cells, measured by flow cytometry, could serve as pre-screening indicators to distinguish between culture-positive and culture-negative samples (Supplementary Figure 7). The analysis of bacterial cell counts (Supplementary Figure 7a) resulted in an AUROC of 0.88 (95% CI: 0.84 - 0.92). A cutoff of 428 cells/µL yielded a positive predictive value (PPV) of 0.96 (95% CI: 0.89 - 0.99) and a negative predictive value (NPV) of 0.69 (95% CI: 0.63 - 0.78). Additionally, the assessment of total DNA yield (Supplementary Figure 7b) achieved an AUROC of 0.85 (95% CI: 0.79 - 0.89), with a PPV of 0.94 (95% CI: 0.88 - 0.97) and an NPV of 0.64 (95% CI: 0.56 - 0.75) at a cutoff of 273 ng (∼34.25 ng/mL). Therefore, flow cytometry and DNA yield can stratify samples into higher- and lower-probability of clinical positivity.

### Flow cytometry data showed increased leukocyte counts in patients with catheters

Plotting the distributions of cell counts obtained by flow cytometry showed no significant difference in leukocyte counts between female and male patients; however, bacterial cell counts were significantly higher in samples from females (Figure 2a). Samples from catheterized patients had significantly higher leukocyte counts than those from non-catheterized patients (Figure 2b). Additionally, bacterial cell counts in catheter urine were higher, although the difference did not appear significant. Bacterial counts across different urine types showed a bimodal distribution. This was most noticeable in non-catheter samples but was present in all sample types, with peaks around 10^4^ and 10^7^ cells/mL. While flow cytometry clearly demonstrates this bimodal trend, it is less apparent in CFU counts from traditional microbiological methods (Figure 2c and 2d).

**Figure 2:**
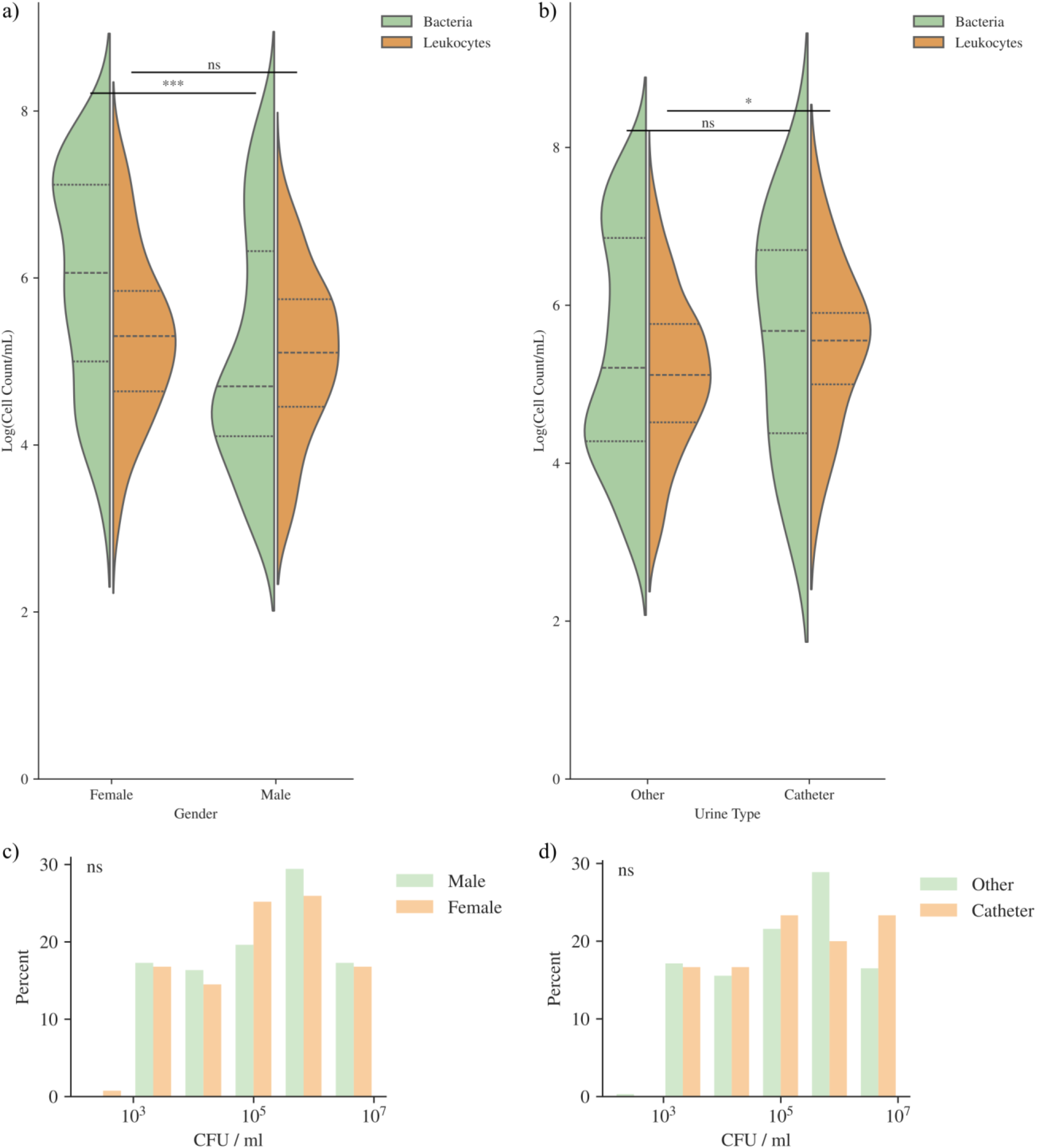
Distributions of flow cytometry cell numbers and CFU stratified by urine type and sex. The distributions of flow cytometry cell numbers per mL (bacteria and leukocytes) are shown as violin plots stratified by (a) patient sex and (b) urine type (catheter urine and other urine). The distributions of colony-forming units (CFU/mL) are shown as histograms stratified by (c) patient sex and (d) urine type. Hypothesis testing was performed using a two-sided Mann-Whitney U test to evaluate whether distributions differ significantly. P-values were corrected for multiple testing using the Bonferroni method. Significance codes: ***: p-value < 0.001, **: p-value < 0.01, *: p-value < 0.05, ns: not significant.

### A significant correlation was observed between the colony-forming units (CFU/mL) and the flow cytometry data

Linear regression analysis of leukocyte and bacterial cell count, and CFU/mL shows a significant (p-value < 0.0005) correlation between the number of leukocytes and the number of bacterial cells measured using flow cytometry (R^2^ = 0.35, SE = 0.05; Supplementary Figure 9a). This trend also significantly (p-value < 0.0005) correlates with CFU/mL counts from conventional microbiological testing, although there is increased variability. Regression of bacterial cell numbers against CFU showed an R^2^ of 0.33 and a standard error of 0.07 (Supplementary Figure 9b). Regression of leukocyte cell numbers against CFU showed an R^2^ of 0.20 and a standard error of 0.06 (Supplementary Figure 9c).

### Genome coverage as a predictor for AST sensitivity

Linear regression and AUROC analysis were used to assess how the breadth of genome coverage relates to the sensitivity of AST predictions on a per-species basis (Supplementary Figure 10). Three of the most abundant uropathogenic species had sufficient data for this analysis. Both *E. coli* and *K. pneumoniae* showed a significant positive correlation between breadth of coverage and AST sensitivity (p-value < 0.005). AUROC analysis suggests a significant number of isolates had AST predictions with >95% sensitivity when coverage was around 20% or higher. The AUROC values were 0.90 and 0.83 for *E. coli* and *K. pneumoniae*, respectively, indicating that genome coverage could potentially be a good predictor of AST sensitivity for these species. However, the *E. faecalis* data revealed no significant correlation between predicted AST sensitivity and genome coverage.

### Different bacterial species demonstrated a range of resistance mechanisms

Species with ARGs detected in at least two samples were plotted in Supplementary Figure 11a, illustrating the relative abundance of resistance mechanisms by type. The most common resistance mechanisms identified were antibiotic efflux, followed by β-lactamases. Efflux-related genes were further divided into ATP-binding cassette (ABC), major facilitator superfamily (MFS), and resistance-nodulation-cell division (RND) type efflux pumps. Most data were available for *E. coli*, where several efflux-related genes from the RND and MFS families were detected, including the *marA* gene, which acts as a global inducer of MFS efflux pumps and down-regulates OmpF porin synthesis. In *E. coli*, class C β-lactamase genes (*CMY* and *EC*) and the *bacA* gene, which confers bacitracin resistance through target modification, were also detected. mNGS detected the ABC-type efflux complex *efrAB* in *E. faecalis*, the MFS-type efflux gene *efmA* in *E. faecium*, two MFS efflux genes, and the *ermT* methyltransferase gene in *S. aureus*, as well as the *tetM* ribosomal protection protein gene in *S. agalactiae*. Class A β-lactamase genes (*CTX-M* and *SHV*) were found in *K. pneumoniae*, along with the RND efflux gene complex *acrAB*. In *P. aeruginosa*, β-lactamase genes of class D (OXA) and class C (PDC) were identified, along with multiple RND family efflux genes.

### The number of virulence genes varies depending on the leukocyte and bacterial cell counts

The changes in the abundance of virulence factors with leukocyte and bacterial cell numbers were examined in Figure 3. Virulence factors were correlated with leukocyte counts (Figure 3a) to determine whether any were associated with increased host immune responses. An increase in adhesion-related virulence factors and a decrease in exotoxins are associated with rising leukocyte counts. Samples with a low leukocyte count (<10/mL) showed a high proportion of virulence factors involved in immune modulation. Interestingly, virulence factors related to immune modulation were more abundant in samples with 10^5–6^ leukocytes/mL than in those with 10^3^ leukocytes/mL. Virulence factors associated with effector delivery systems were most prevalent in samples with the lowest leukocyte counts and least prevalent in samples with 10^3^ leukocytes/mL. Biofilm formation factors were most abundant in samples with the lowest leukocytes and least in those with 10^4^ leukocytes/mL. Additionally, virulence factors changed with bacterial cell numbers (Figure 3b). At low bacterial counts, exotoxins, immune modulation, and adherence are the most common categories of virulence factors. Also, immune modulation, biofilm formation, and effector-delivery-system-related virulence factors were observed at 10^4^ to 10^6^ leukocytes/mL. This indicates that as bacterial population density increases, the relative abundance of adherence factors, nutritional and metabolic factors, and effector delivery systems also increases.

**Figure 3:**
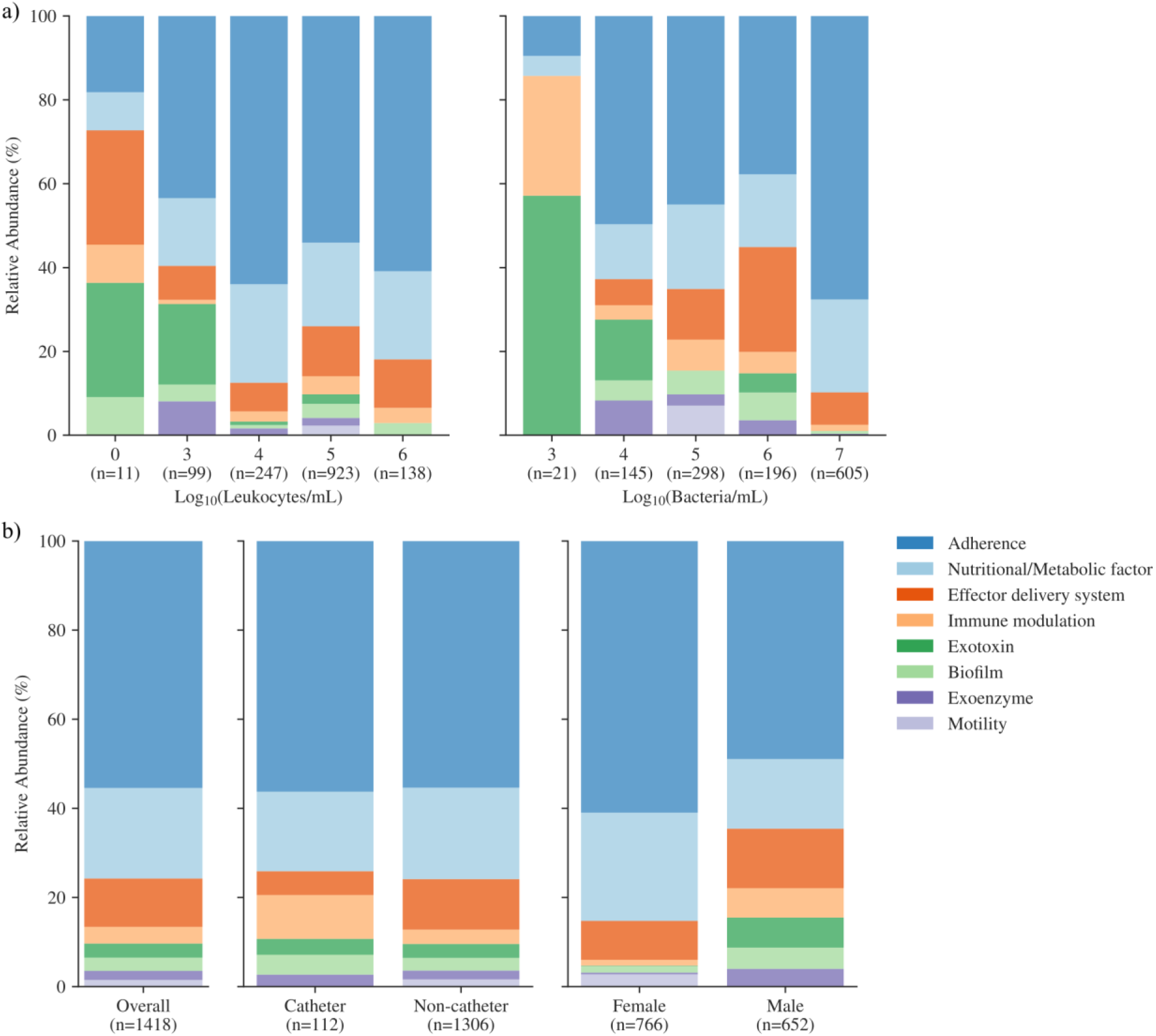
The relative abundance of virulence factors identified through mNGS. Virulence factors are stratified in various ways. Subfigure (a) shows virulence factors binned by the base-10 logarithm of leukocyte and bacterial cell numbers measured using flow cytometry. Subfigure (b) shows the relative abundance of virulence factors overall, stratified by catheterization status and by gender. The total number of virulence genes included is shown (n).

At the species level, virulence factors identified by mNGS also showed variability across different species. Supplementary Figure 11b illustrates the differences in the relative abundance of virulence mechanisms by type for relevant uropathogenic species. Among the various types of virulence factors, adherence (∼60%) and nutritional/metabolic factors (∼20%) were the most common in *E. coli* and *K. pneumoniae*. These were followed by exoenzyme and other virulence factors. In contrast, there was significant variation within the *Enterococci* group, with *E. faecalis* displaying a range of virulence factors, primarily adherence (∼50%) and nutritional/metabolic factors (∼20%). Conversely, *Enterococcus faecium* primarily possessed an effector-delivery system virulence (∼70%), with adherence and nutritional/metabolic factors also accounting for substantial proportions of the identified genes. *P. aeruginosa* showed diverse levels of abundance across different virulence gene classes. Motility-related virulence factors were found in *E. faecalis* and *P. aeruginosa* but were absent from other species. Unlike other species, *S. aureus* and *S. agalactiae* contained abundant exoenzyme- and exotoxin-related virulence genes. The presence of immune-modulation-related virulence factors was significantly higher in *E. faecalis* and *S. agalactiae* compared to other species.

### Diverse co-occurrences among the uropathogens

From the entire cohort from phases I and II, 91 samples were identified as polymicrobial (see Supplementary Figure 1). The most common uropathogens co-occurring in many polymicrobial samples were *E. coli*, *E. faecalis*, and *K. pneumoniae* (Figure 5a). Specifically, *E. coli* was found alongside *E. faecalis* in 16 samples and with *K. pneumoniae* in 3 samples. Additionally, *E. faecalis*, excluding *E. coli*, co-occurred with *K. pneumoniae* in 5 samples, *Morganella morganii* in 6 samples, *P. aeruginosa* in 4 samples, and *S. aureus* in 5 samples. Along with common uropathogens, other urobiome bacteria, including *Actinotignum schaalii*, *Aerococcus urinae*, *Aerococcus sanguinicola*, and *Streptococcus constellatus*, were identified. These bacteria coexisted with dominant uropathogens, including *E. coli*, *E. faecalis*, and *K. pneumoniae*. Pathogens with multidrug resistance most frequently showed concurrent resistance to β-lactam antibiotics, with ampicillin and amoxicillin (42 cases), ampicillin and amoxicillin/clavulanic acid (54), ampicillin and cefuroxime (37), amoxicillin and cefuroxime (14), and ampicillin and cefpodoxime (25) (Figure 5b).

**Figure 4:**
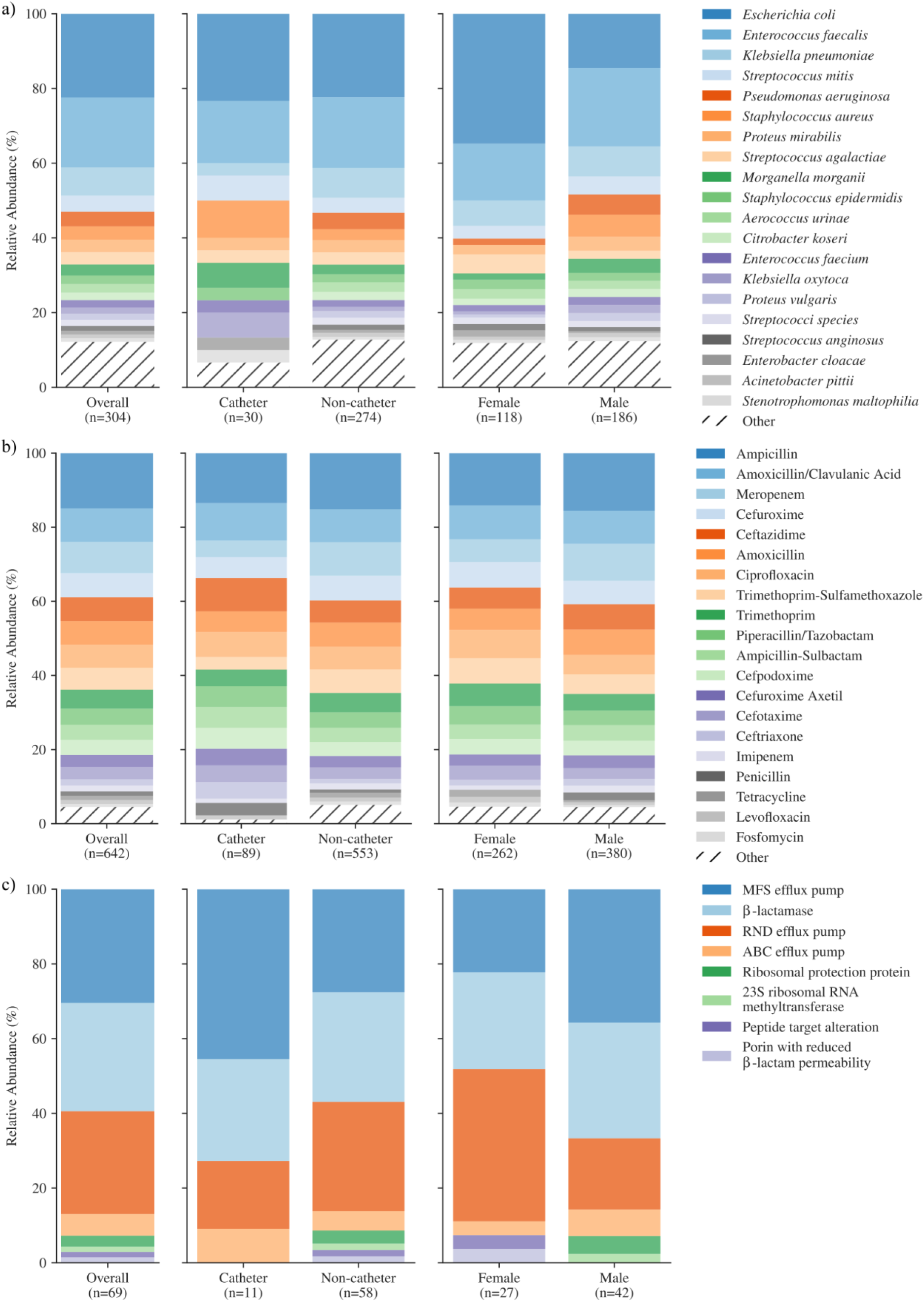
Risk factors (pathogen, AST, ARGs). The relative abundance of the top 20 (a) identified pathogens (MALDI-TOF), (b) Antibiotic resistance (routine AST), and (c) resistance mechanisms are shown overall and stratified by catheterization status and gender.

**Figure 5:**
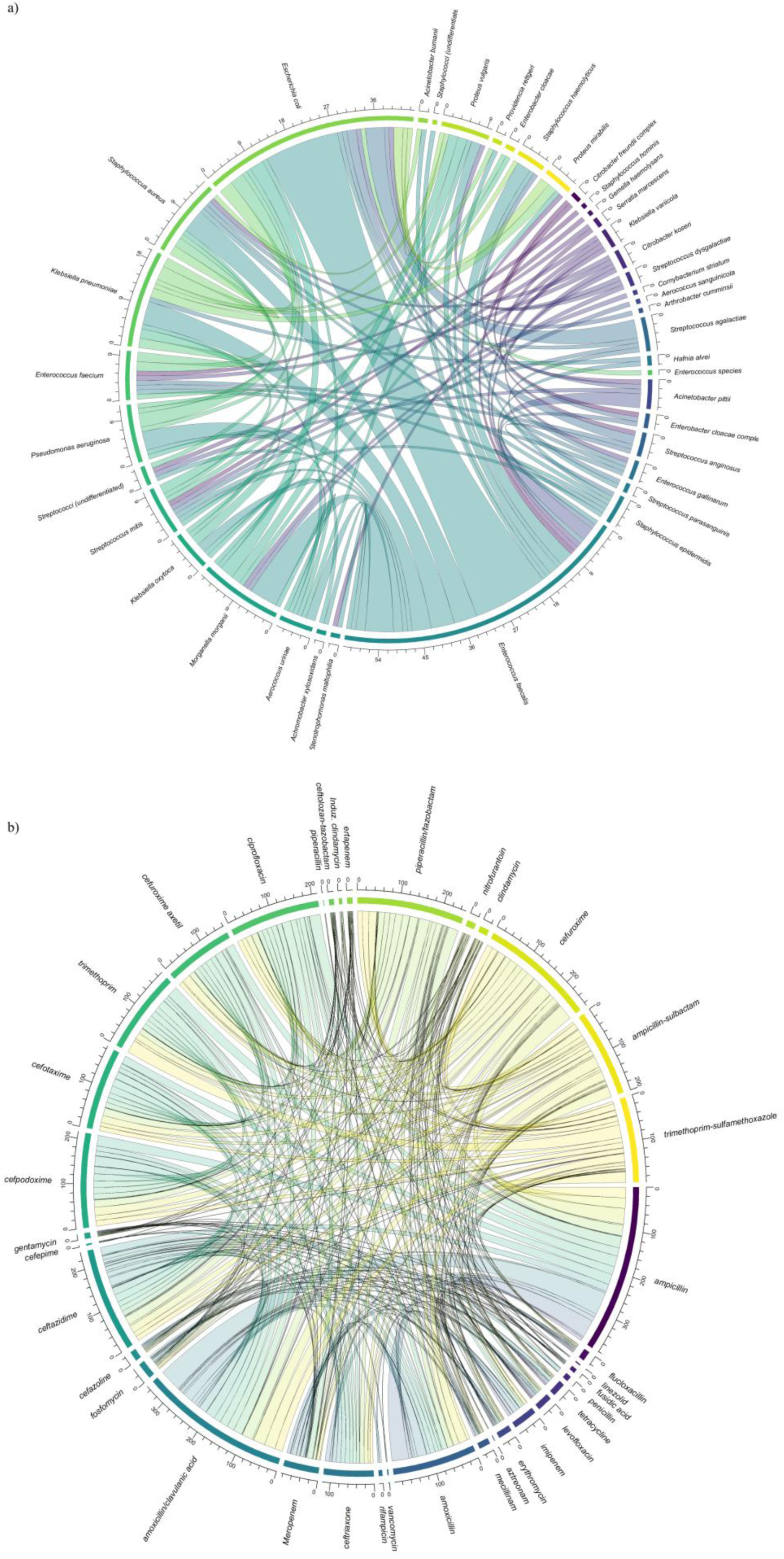
Co-occurrence of pathogens and antibiotic resistance. Subfigure (a) shows a chord diagram depicting the co-occurrence of pathogens in polymicrobial samples processed via the URINN method. Each species sector is annotated with absolute numbers to clarify the number of samples in which each species occurs. The size of the connections between sectors (the ribbon’s thickness) reflects how frequently the connected species appear in a common sample. The subfigure (b) shows a chord diagram depicting the co-occurrence of resistance in samples processed via the URINN method. Each antibiotic sector is annotated with absolute numbers to clarify the number of samples in which resistance to each antibiotic occurs. The size of the connections between sectors (the ribbon’s thickness) reflects how frequently resistance to the connected antibiotics appears in a common sample. Colors are used to visually distinguish these connections.

### Characterization based on the risk factors: Catheter and gender

When differentiating among catheter and other urine sample types beyond *E. coli* and *E. faecalis*, *S. aureus*, *Morganella morganii*, and *Klebsiella oxytoca* were more prevalent in catheterized urine samples than in non-catheter samples. In non-catheter urine samples, *K. pneumoniae* and *P. aeruginosa* were more abundant (Figure 4a). Analysis of virulence factors in catheterized and non-catheterized samples showed that those associated with immune modulation and biofilm formation were more prevalent in catheter-derived samples. Conversely, non-catheter urine samples contained higher levels of effector delivery system-related factors and virulent factors linked to mobility (Figure 3b). Catheter urine samples also exhibited a greater proportion of resistance to cephalosporins, including cefotaxime, cefpodoxime, ceftazidime, ceftriaxone, and cefuroxime, while other urine samples demonstrated a high level of resistance to meropenem, primarily due to intrinsic resistance in *Enterococcus* species (Figure 4b). The most common resistance mechanism in catheter-derived samples was the OXY β-lactamase gene, which is found in *K. oxytoca*. The β-lactamase *EC-5* was also detected in an Escherichia coli from a catheter urine sample. Additional resistance mechanisms in catheter urine samples involved antibiotic efflux systems, including *acrAB-TolC* and *efrAB*. In non-catheter urine samples, the β-lactamase genes *SHV* and *CTX-M* were identified in *K. pneumoniae* and *Proteus mirabilis*, and *blaACT* was found in *E. cloacae* and *P. aeruginosa* (Figure 4c).

A clear difference in the abundance of uropathogens was observed based on gender. *E. coli*, *E. faecalis,* and *K. pneumoniae* were the most common uropathogens in both genders. However, *E. coli* was more prevalent in female patients than in male patients. In addition, *P. aeruginosa*, *S. aureus,* and *M. morganii* are more abundant in males, whereas *Streptococcus agalactiae* is more abundant in females (Figure 4a). There is also a clear difference in virulence factors between females and males (Figure 3b). Adherence and nutritional/metabolic factors were more common in females, while virulence factors related to exotoxins, exoenzymes, effector delivery systems, biofilm formation, and immune modulation were more prevalent in males. Virulence factors linked to motility were absent in male samples. Females showed a slightly higher rate of resistance to fluoroquinolones and tetracyclines, whereas males had a higher rate of resistance to penicillin and carbapenem antibiotics (Figure 4b). In female samples, the following β-lactamase genes were identified: *CMY* (E. coli), *OXA* (*P. aeruginosa*), *OXY* (*K. oxytoca*), and *PDC* (*P. aeruginosa*). In males, the β-lactamase genes *DHA* (*M. morganii*), *SHV* (*K. pneumoniae*), and *ACT* (*Enterobacter cloacae*) were found (Figure 4c).

## Discussion

Currently, bacterial culture is the standard method for diagnosing UTIs, but it has limitations, including prolonged turnaround times and lower detection sensitivity^14,15^. This study expands our previously developed mNGS method^13^, URINN (formerly known as the “optimized method”), by enhancing and testing it on a much larger cohort of 296 patient urine samples. This method’s performance was compared with clinical microbiology results obtained with the MALDI-TOF and VITEK-2 systems. The findings show that, despite a significantly larger dataset of 296 samples compared with 50 in phase 1, pathogen identification accuracy remains nearly unchanged. Furthermore, susceptibility prediction accuracy (92%) and specificity (99%) have improved slightly compared with those in our previous proof-of-concept study^13^. The method has a low detection limit, enabling identification of pathogens at 10^3^ CFU/mL with high precision. We also emphasize the benefits of metagenomic sequencing by analyzing pathogenic traits in both phase I and phase II cohorts, extending beyond mere taxonomy and antibiotic susceptibility. The analysis identified *E. coli*, *E. faecalis*, and *K. pneumoniae* as the leading uropathogens, each exhibiting a variety of virulence factors, primarily related to adhesion and nutrition. These pathogens often co-occur with other uropathogens in polymicrobial samples.

The phase II cohort includes 296 urine samples from male and female patients collected routinely at the University Hospital in Giessen. The cohort exhibits a complex microbiological profile, with 50% (149/296) of samples being culture-positive. Among these, 58% (87/149) were monomicrobial, while 42% (62/149) were polymicrobial. Of 232 bacterial pathogens identified, routine AST was performed on 163 (Supplementary Table 4), of which 78% (132/163) were resistant to at least one antibiotic. Among the resistant pathogens, 41% (67/143) were multidrug-resistant, i.e., resistant to three or more antibiotics.

Minor modifications were made to the optimized phase I protocol to improve performance, considering the diverse nature of the urine samples. In the host-depletion step, the saponin concentration was increased from 2.2% to 3%. A subsequent qPCR assay did not reveal any significant differences in host depletion levels. This is likely due to the fixed number of WBCs spiked into the samples. However, host-level variation among clinical samples may lead to more effective host depletion at higher saponin concentrations. Additionally, the results show that this does not affect bacterial DNA, consistent with our previous study, in which we tested different saponin concentrations (2-5%) and observed only minor variations in the planktonic growth rates of *E. coli* and *S. aureus*^16^. During DNA extraction, the lysis steps were modified to reduce DNA fragmentation and enhance extraction from a urobiome perspective. Additionally, ice cooling was added to prevent thermal or nuclease-related degradation of nucleic acids during bead-beating. Through URINN, we also detected Candida sp. in seven clinical samples (Supplementary Table 6), demonstrating the method’s potential to identify fungi alongside bacteria. Although primarily designed for bacterial pathogens, these findings suggest it could be modified for comprehensive urobiome metagenomics.

The URINN method demonstrated a pathogen identification accuracy of 92% (257/279), with a sensitivity of 97% (199/206), specificity of 79% (58/73), precision of 93% (199/214), and an NPV of 89% (58/65). Despite involving a larger cohort, these results closely align with those from the optimized method in our proof-of-concept study (Table 1) and exceed the performance reported in similar clinical studies of UTI samples^17–20^. The apparent decrease in specificity results from the inability to confirm the presence of additional pathogens identified through other independent methods. Metagenomics provides in-depth insights into microbiomes, including virulence factors, resistance mechanisms, the presence of species with unknown clinical significance, commensal flora, and potential false positives from environmental or other contaminants. The URINN metagenomics method can identify species that traditional tests might miss, raising questions about how labs should communicate these results to clinicians and patients^21^. Protocols should be established to identify clinically relevant findings and to ensure clear communication with microbiologists, clinicians, and patients. Here, we report the pathogen identification results in accordance with the guidelines of Johnson et al. (2025)^21^ and have classified the identified bacteria as pathogenic, possibly pathogenic, or benign (Supplementary Tables 6 & 7).

Antibiotic resistance prediction from mNGS data is perhaps the most challenging aspect, as the presence of the gene does not necessarily indicate resistance^22^. For antibiotic susceptibility prediction, the method achieved 92% (1330/1446) accuracy, with 64% (177/277) sensitivity, 99% (1153/1169) specificity, 92% (177/193) precision, and a NPV of 92% (1153/1253), consistent with previous proof-of-concept results (Table 1). Reporting of antibiotic resistance genes follows established conventional AST panels at the per-pathogen level, while also considering the intrinsic resistance of these species. Most current mNGS studies generate susceptibility predictions based on antibiotic classes^18,23^, which limits their clinical applicability. This study further improves the high specificity of our previously reported AST prediction method, demonstrating that it does not overpredict resistance and is less likely to lead to unnecessary antibiotic use. Combining results from both study phases results in an accuracy of 91% (1919/2099) and a specificity of 98% (1660/1701) for AST concordance across 2099 antibiotics, emphasizing the robustness of the method.

However, the method’s AST prediction sensitivity is relatively low, with both phases combined achieving 65%. This is mainly because it performs poorly at predicting resistance mechanisms to trimethoprim and ciprofloxacin (Supplementary Figure 4). Resistance to these drugs often results from chromosomal mutations, overexpression of efflux pumps, or plasmid-mediated resistance^24^. Interestingly, a study of uncataloged genetic variations in antimicrobial resistance gene families in *E. coli* found that 3.4% of ciprofloxacin resistance and 12.7% of trimethoprim resistance could not be explained by any known antibiotic resistance genes^25^. In comparison, the unexplained resistance in trimethoprim is much higher, as both *dfrA* (>30 known allele variants^26^) and *folA* are poorly characterized. Prediction of resistance to amoxicillin-clavulanic acid and ampicillin was also low, mainly in *E. coli*. Unlike other antibiotic resistances, resistance to amoxicillin-clavulanic acid (AMC) generally does not stem from a single ARG but rather from a combination of mechanisms, such as promoter mutations that cause hyperproduction of ampC, overexpression or inhibitor-resistant TEM variants, and decreased permeability, making it difficult to predict^27^. A large phenotypic and genotypic study of 234 *E. coli* isolates identified 69% categorical agreement for AMC, the lowest among the 11 tested antibiotics^28^. Additionally, low genome coverage can lead to underreporting of resistance in metagenomics^29^.

We also explored the link between genome coverage breadth and the sensitivity of AST predictions for the three most abundant species, *E. coli*, *E. faecalis*, and *K. pneumoniae*. AUROC analysis indicated that a substantial number of isolates had AST predictions with >95% sensitivity at approximately 20% coverage. This suggests that having broad genome coverage might be necessary to achieve high-quality prediction of AST sensitivity in these species. However, for *E. faecalis*, no significant correlation was observed between the breadth of genome coverage and the accuracy of AST sensitivity prediction. This indicates that increasing coverage does not improve AST prediction performance. This could be due to a lack of well-characterized resistance mechanisms in *E. faecalis*^30^ beyond vancomycin resistance^31^. To our knowledge, this is the first study to explore this correlation, and additional data, along with follow-up research, are needed to establish threshold values based on genome coverage for evaluating the accuracy of AST predictions.

Analysis of flow cytometry data showed a significant linear correlation between leukocyte and bacterial cell counts (Supplementary Figure 9). This indicates that higher pathogen levels can trigger a stronger immune response. However, considerable variability in leukocyte numbers remains that cannot be fully explained by bacterial abundance alone (R2 = 0.35, standard error = 0.05), likely due to underlying comorbidities in the patient group, which can also trigger immune responses. A similar trend is observed with leukocytes and bacterial cell counts relative to CFU/mL data (R2 = 0.33 and 0.20, respectively), but biological variability remains. While flow cytometry has been used to detect significant bacteriuria^32^, the difference between cell counts and CFU/mL data arises from flow cytometry detecting all cell types (live, dead, viable but non-culturable, aggregates, etc.)^33^, whereas culture measures actively surviving and culturable cells. AUROC analysis of DNA yield and flow cytometry identified cutoff points of 428 cells/µL and 34.25 ng/mL (or 273 ng for 8 mL). These values align with those reported in our previous study. Notably, applying URINN to a larger sample set produced similar cutoffs. However, further research is needed before these can be used as screening criteria for culture positivity.

Virulence factors are essential for pathogenicity because they enable pathogens to infect hosts, promote colonization, and evade immune responses. These factors also offer valuable insights into the pathogen and can assist in assessing the prognosis of UTIs. Overall, within the current cohort, adherence virulence factors were the most common among the factors studied, a pattern consistent across samples grouped by catheter type and gender (Figure 3). This trend is also observed when samples are grouped by the most prevalent uropathogens in the cohort, including *E. coli*, *E. faecalis*, and *K. pneumoniae* (Supplementary Figure 11b), which display similar abundances of VF, primarily related to adherence, nutritional and metabolic support, and effector delivery systems. This is expected because adherence is a key initial step in UTI development and, along with other factors such as nutrition and metabolism, is essential for initiating UTI pathogenesis and successful colonization^34^. Among the adherence group, the most abundant were *E. coli* type 1 fimbriae (*Fim* operon), S fimbriae (*Sfa* cluster), and F1C fimbriae (*Foc* cluster)^35^. Uropathogenic *E. coli* (UPEC) harbor significantly more fimbrial genes than fecal or commensal pathotypes^36,37^. These factors were identified in various midstream and catheter samples. These virulence genes have been shown to facilitate UPEC adherence to the bladder epithelium, invade, transform mast cells into caveolae, and evade the host immune response^38,39^, thereby promoting colonization and persistence^40^. This bacterial ability to invade host cells is closely linked to recurrent UTIs (rUTIs)^41^. Similarly, virulence genes linked to siderophores such as aerobactin, enterobactin, and yersiniabactin were also identified. These nutritional and metabolic factors aid *E. coli* (aerobactin and yersiniabactin)^34,42^ and *K. pneumoniae* (aerobactin and enterobactin)^43^ in scavenging iron from the bladder environment. A study by Lei *et al*. identified these nutritional and metabolic virulence factors as among the most strongly associated with rUTI relapse and remission^44^. All prevalent uropathogens also possess additional virulence genes related to immune modulation and biofilm formation, although the proportion is comparatively higher in *K. pneumoniae* and *E. faecalis*. This is expected, as the *E. faecalis* identified here harbors *esp* and *ebp,* which are known to form biofilms in urinary catheters^34^ and are often detected in polymicrobial infections^45^. Additionally, *E. faecalis gelE-sprE* exoenzyme genes were found in 5 out of 18 samples. These genes have previously been identified as important biomarkers of biofilm strength^46^.

mNGS identified a higher proportion of virulence factors associated with biofilm formation in *P. aeruginosa* compared to *E. coli* and *K. pneumoniae*. *P. aeruginosa* is known to form biofilms to resist antibiotics and environmental stresses. As a result, it is more likely than *E. coli* and *K. pneumoniae* to develop biofilms on medical devices^47^. However, the analyzed cohort did not show an increased prevalence of P. aeruginosa among catheterized patients, potentially due to the relatively small sample size. The high proportion of exotoxin-related virulence factors in S. aureus is expected, as it produces various exotoxins and proteases that facilitate host colonization and contribute to its pathogenicity^48^. *S. aureus* was more common in patients with catheters, which also accounts for the approximately 20% presence of biofilm-related virulence genes (Supplementary Figure 11b). It has been noted that *S. aureus* utilizes fibrinogen released during catheter-induced inflammation to enhance its virulence and promote infection, explaining its higher prevalence in catheterized patients and the increased risk of bacteremia^49^. Most virulence factors identified in *Streptococcus agalactia*e were associated with the β-hemolysin/cytolysin exotoxin (*cyl gene*), which is known to promote bladder colonization and inflammation in UTI^50^. These virulence factors, in combination with capsular sialic acid, augment *E. coli* survival in the bladder when co-infection occurs^50^ and can lead to persistent and recurrent infections. Therefore, the above-listed virulence factors enable uropathogens to persist in the urinary tract and cause recurrent infections^41^.

Numerous species- and strain-specific virulence factors can influence the risk of rUTI, with infection severity typically reflecting the balance between the host’s defenses and the pathogen’s virulence^51^. However, only a few studies have examined the connection between VFs and UTI severity, recurrence, and persistence. mNGS could support this research, potentially leading to the development of vaccines targeting key virulence factors and enabling more personalized treatments^52^. For example, co-administration of mannosides has been shown to inhibit intestinal colonization of UPEC by blocking FimH-mediated adhesion, thereby preserving the microbiota and reducing UTI recurrence and incidence^53^. We also examined the relationship between leukocyte counts and virulence factors (Figure 3a). An increased presence of adherence, nutritional/metabolic, and immunomodulation factors, which are positively associated with leukocyturia, indicates potential bacterial colonization^54^. However, caution should be taken when interpreting this, as the immune response is affected by host factors.

Additionally, we analyzed the abundance of uropathogens, their antibiotic resistance profiles, and associated resistance mechanisms (Figure 4), focusing on two risk factors: catheterization and gender. Both groups—catheter and non-catheter—included all traditional uropathogens. However, the catheter group showed higher abundances of *M. morganii*, *K. oxytoca*, and *S. aureus*, which have also been frequently observed in other studies of catheter-associated infections^55,56^. Meanwhile, the midstream void samples exhibited a more diverse uropathogen profile, likely reflecting microbial communities from the bladder and lower urinary tract^57^. When grouped by gender, the relative abundance of *E. coli* was higher in female samples, whereas the abundance of non *E. coli* was higher in male samples*. coli* pathogens were greater in male samples. The gender-based prevalence of the uropathogens likely reflects the combination of anatomical and clinical factors^58^. Analyzing resistance mechanisms across different risk factor groups reveals that ARGs are distributed among various functional categories. This suggests a likely multifactorial resistance system in which resistance results from the combined effects of multiple ARGs rather than from a single dominant gene. Although the data do not conclusively demonstrate polygenic resistance, the identification of multiple resistance-related mechanisms supports earlier studies suggesting that resistance phenotypes can emerge from the collective influence of diverse genetic factors^59^.

It is well known that cases of UTI with systemic involvement, often referred to as complicated UTI, are often polymicrobial, particularly when risk factors such as advanced age, a weakened immune system, and comorbidities are involved^60^. This observation also applies to the current cohort, in which 42% of samples are polymicrobial (Supplementary Figure 1). It is also crucial to further investigate pathogen co-occurrence among these populations, as polymicrobial bacteraemic UTIs are often associated with higher morbidity than UTIs caused by a single pathogen^61^. In the current cohort, *E. coli* and *E. faecalis* frequently co-occur across samples (Figure 5a). *E. faecalis* has often been reported to be part of the polymicrobial community^62,63^. Additional studies have shown that *E. faecalis* can modulate the host response and promote the virulence of the co-infecting species, such as *E. coli*^64,65^. A retrospective multicenter analysis of 99415 male outpatient samples in Germany from 2015 to 2020 identified *E. coli* and *E. faecalis* as the leading co-infecting pathogens, accounting for 48.4% and 46.8% of all polymicrobial infections, respectively^66^. Moreover, the presence of *E. coli* and *E. faecalis* is associated with recurrent UTI, particularly in women^67^.

mNGS, with its agnostic approach, can also provide insights into the urobiome. Although there is no consensus on what characterizes a healthy microbiome in males and females, changes in the microbiome have been linked to various diseases^68^. In the current cohort, additional commensal/opportunistic pathogens were identified among the samples (Supplementary Table 7). Species such as *A. urinae and A. schaalii* are increasingly associated with UTIs, especially in patients with underlying health conditions^69^. Furthermore, other microbiome species are known to impact host susceptibility to UTIs. For example, women with overgrowth of anaerobic species, such as Gardnerella vaginalis (also found in this cohort), are more likely to develop UTIs than those with a more diverse microbiota, such as *Lactobacillus*^70^. Consequently, research on the urobiome is expanding to include cases like prostate cancer^71^, urinary incontinence^68^, and kidney transplants^72^, all of which have been linked to urobiome dysbiosis. This questions the traditional UTI treatment approach rooted in Koch’s postulates^73^, which links pathogenicity to a single organism and highlights the growing importance of the urobiome in managing UTIs.

A limitation of the current study that needs to be addressed to improve metagenomic sequencing is the use of whole-genome amplification for samples with low DNA yields. These samples typically have low bacterial biomass, around 10^3^ CFU/mL, which is also the detection limit (LOD) of the URINN method. It has previously been reported that the WGA could generate species-specific bias^74^. This is less consequential, as 68% (26/38) of amplified samples were monomicrobial (Supplementary Tables 1 & 2). Therefore, additional evaluation is necessary to optimize the WGA method and minimize any potential bias.

In conclusion, this study demonstrates the successful optimization and large-scale validation of the URINN metagenomic sequencing workflow for rapid and accurate pathogen and antimicrobial susceptibility profiling in complicated UTIs. The method achieves over 99% accuracy in identifying pathogens at the sample level and 99% specificity in predicting antibiotic susceptibility, thereby minimizing overprediction. Its detection limit of 10^3^ CFU/mL is clinically relevant for potential UTI diagnosis. It can also detect *Candida spp* in clinical samples, demonstrating its potential for pangenome applications. Beyond pathogen and AST detection, URINN provides comprehensive insights into antimicrobial resistance mechanisms, virulence factors, and polymicrobial interactions that are often overlooked in traditional diagnostics. Stratification by clinical risk factors, including gender and catheterization status, further revealed distinct microbial and resistance profiles that could inform more personalized treatment strategies. There was no significant difference in leukocyte counts between genders, but bacterial load was significantly higher in female patients overall. Differences were also observed between catheterized and non-catheterized patients. The catheterized group had significantly higher leukocyte counts. Additionally, catheterized urine samples showed a higher rate of cephalosporin resistance. Insights into these risk factors can facilitate more tailored development of treatment protocols and improve the clinical management of UTIs.

## Methods

### Study design

In our previous proof-of-concept study, we developed and evaluated 11 methods—8 in-house and 3 commercial—using 78 clinical samples, and identified the optimized method as the most effective. Building on this, we made minor adjustments to the optimized method and subsequently tested it on a cohort of 296 UTI patient samples (Supplementary Tables 1 and 2). The improved method will be referred to as the URINN method.

### Fine-tuning of the optimized method

In the current study, the URINN method was used to selectively remove host material and extract bacterial DNA. The protocol was applied with minor modifications: instead of 2.2% saponin, as used in the optimized method, the concentration was increased to 3%, and the volume of M-SAN was raised to 20 µL. Next, 400 µL of cold TE buffer was added to the pellet, which was then transferred to the bead-beating step. This was followed by treatment with 260 µL Naxtra lysis buffer and 5 µL of proteinase K. Finally, the NAxtra™ Blood magnetic beads were replaced with those of NAxtra™ 2.0 Blood total nucleic acid extraction kit (LSNXD096, Lybe Scientific AS, Trondheim, Norway).

### Evaluating the effect of 2.2 and 3% saponin concentration and qPCR assay

Spiked urine samples were utilized to assess the impact of two saponin concentrations on host depletion and microbial DNA. Briefly, clinical conditions were simulated by spiking urine samples from healthy donors, confirmed by dipstick analysis, with white blood cells isolated from the buffy coat at 1 × 10^4^ cells/mL. Additionally, the samples were inoculated with *E. faecalis* at a final concentration of 10^5^ CFU/mL^13^. DNA extraction was then carried out using the URINN protocol at 2.2% and 3%, with all extractions performed in triplicate (n=3) qPCR assays were conducted to assess the impact of increasing saponin concentrations on host and bacterial DNA extraction, targeting the human β-actin gene and the *UspA* gene, as previously described^13^. Each qPCR assay contained 3 μL of 5X HOT FIREPol® EvaGreen® Supermix (Solis BioDyne, Estonia), 0.2 μM of forward and reverse primers, 10.4 μL of nuclease-free water, and 1 μL of DNA. Assays were performed on a 7500 Fast Real-Time PCR system with an initial denaturation at 95 °C for 720 s, followed by 40 cycles at 95 °C for 25 s, 60 °C for 45 s, and 72 °C for 60 s.

### Whole genome amplification (WGA)

Routine culture-positive samples with DNA concentrations below 200 ng after purification with AmpureXP were amplified using the EquiPhi29 DNA amplification kit. Before amplification, the purification was performed using AMPure XP beads at 1.0X concentration. The beads were added to the samples and mixed with a Hula Mixer (VWR:10136-084) for 10 minutes, then pelleted on a DynaMag-2 rack (Thermo Fisher Cat. No. 12321D). The resulting pellet was washed twice with 1 mL of 80% ethanol. Finally, the DNA was eluted in 15 µL of nuclease-free water. The EquiPhi29 DNA amplification kit (Thermo Fisher, Cat. No. A65393) was used according to the manufacturer’s instructions, with the following adjustments: 12 µL of the sample was used for amplification, and the amplification was performed at 42°C for 60 minutes. After the WGA, the samples were debranched by adding 10 U of S1 nuclease (Thermo Fisher, Cat. no. EN0321) to 5X S1 nuclease reaction buffer, bringing the final concentration to 1X. For debranching, the samples were incubated at 21°C for 2 minutes, then inactivated by adding 2 µL of 0.5 M EDTA and incubating at 70°C for 10 minutes. This was followed by an AMPure XP purification at 1.0X bead concentration, after which the samples were used for library preparation.

### Sampling criterion and screening using flow cytometry data

A total of 296 patient urine samples were prospectively collected from patients with UTIs. As in our previous study, the cohort included samples that were positive for leukocytes or for leukocytes and nitrates. In addition, we applied a bacterial cell count cutoff based on the flow cytometry thresholds reported in the previous study (see the first paper). A cutoff of 251 bacterial cells/uL was used as the flow cytometry screening criterion for clinical samples. Since the earlier cutoffs were derived from a smaller dataset, samples excluded from flow cytometry were cross-checked for culture positivity without revealing any pathogen information. Any positive samples that were missed were then extracted and sequenced, and the results were correlated with clinical data. The control cohort samples were obtained from patients without symptoms who presented for a preventive medical examination in the urology ward. All the samples in this cohort were culture-negative.

### Routine clinical culture for identifying pathogens and testing antibiotic susceptibility

The routine microbiological analysis of the collected samples, including pathogen identification and antibiotic susceptibility testing, was conducted in accordance with the regulations of the university hospital’s microbiological laboratory, using MALDI-TOF and VITEK-2 systems as previously described^13^. The results were interpreted in accordance with the guidelines established by the European Committee on Antimicrobial Susceptibility Testing (EUCAST).

### Nanopore sequencing of the extracted/amplified DNA samples

The extracted/amplified DNA, post-purification, was sequenced on a PromethION 2 solo device. The library preparation was performed using the Rapid barcoding kit SQKRBK114-96 (Oxford Nanopore, UK) and sequenced on PromethION flow cells (R10.4.1 FLO-PRO114M). All samples sequenced in the study were divided into batches of 48-60 and sequenced on PromethION flow cells.

### Bioinformatics analysis of sequencing data

Data analysis for the detection of pathogens, ARGs, and virulence genes was performed as previously described^13^. Briefly, unclassified reads were recovered using MysteryMaster^75^, reads were mapped to a custom uropathogen reference database using BLASTn, and non-aligned reads were mapped against the NCBI RefProk database using BLASTn with the previously described parameters. Bacterial species were categorized into three groups according to Johnson et al., pathogenic, possibly pathogenic, and benign^21^. Any bacteria classified as uropathogens in the reference database are marked as pathogenic when reads longer than 1 kb are present. Species that could be opportunistic pathogens in UTI (refer to Supplementary Table 7) were considered possibly pathogenic if their abundance was over 50% of all bacterial reads. All other bacterial species that did not meet either criterion but had more than 5% read abundance were classified as benign. After identifying pathogens using BLASTn, reference-based alignments were made using minimap2^76^ and used for the identification of resistance and virulence genes. Abricate^77^ was used with the CARD^78^ and VFDB^79^ databases to identify antibiotic resistance genes and virulence factors, respectively. An additional exclusion criterion was introduced for amplified samples: species with less than 10% genome coverage were excluded from further analysis, regardless of read count.

Linear regression was used to determine the correlation between AST prediction specificity and the breadth of genome coverage. P-values were calculated using the Wald Test with a t-distribution. AUROC analysis was used as described^13^ to investigate whether genome coverage is a predictive measure of high AST specificity. Pathogens were considered to have high specificity (positive) when AST prediction specificity was at least 95%. This analysis was limited to species for which at least 10 individual bacteria had both sequencing data and routine AST data available.

### Statistical analysis, scoring metrics, and limit of detection (LOD)

As previously described^13^, the area under the receiver operator characteristic curve (AUROC) calculations were performed using routine microbiological data as a reference for total DNA yield and the number of bacterial cells from flow cytometry. The accuracy scores for pathogen identification and antibiotic susceptibility prediction were similarly calculated at three levels: sample, pathogen, and antibiotic. The LOD here was defined as the concentration of colony-forming units (CFU) that yields a 95% probability of detecting the pathogen in a sample. The 95% LOD was calculated using a beta-distribution approximation.

### Ethics

The study was approved ethically by the Ethics Committee of Justus Liebig University Giessen, Faculty of Medicine (AZ 158/20) for UTI samples, and by the regional medical and health research ethics committees (REC Helse Sør-Øst) (691177) for healthy donors. The study had a waiver of consent for the collection and use of these samples.

### Data availability

The sequencing data generated in this study are deposited in the European Nucleotide Archive database with the accession number PRJEB107316.

## Acknowledgements

A.B.B., S.B., and R.A. were supported by the Research Council of Norway through the projects OH-AMR-Diag (project number 336420) and UTI-Diag (project number 352514). FW and TH are members of the DFG (German Research Foundation) funded research group BARICADE (FOR5427/1-466687329) and members of the DZIF (German Center for Infection Research; site: Giessen–Marburg–Langen).

## Author contributions

R.A., F.W., T.H., C.I., and T.B.J. planned the study. A.B.B., S.B., and R.A. designed the experiments. A.B.B., I.K., performed the experimental work in discussions with R.A. A.B.B., S.B., and R.A. wrote the manuscript. All authors contributed to the article and approved the submitted version.

