## Supplementary Figure 1 for "Rapid clinical metagenomics enables early tailored therapy in complicated urinary tract infections and strengthens antimicrobial stewardship"

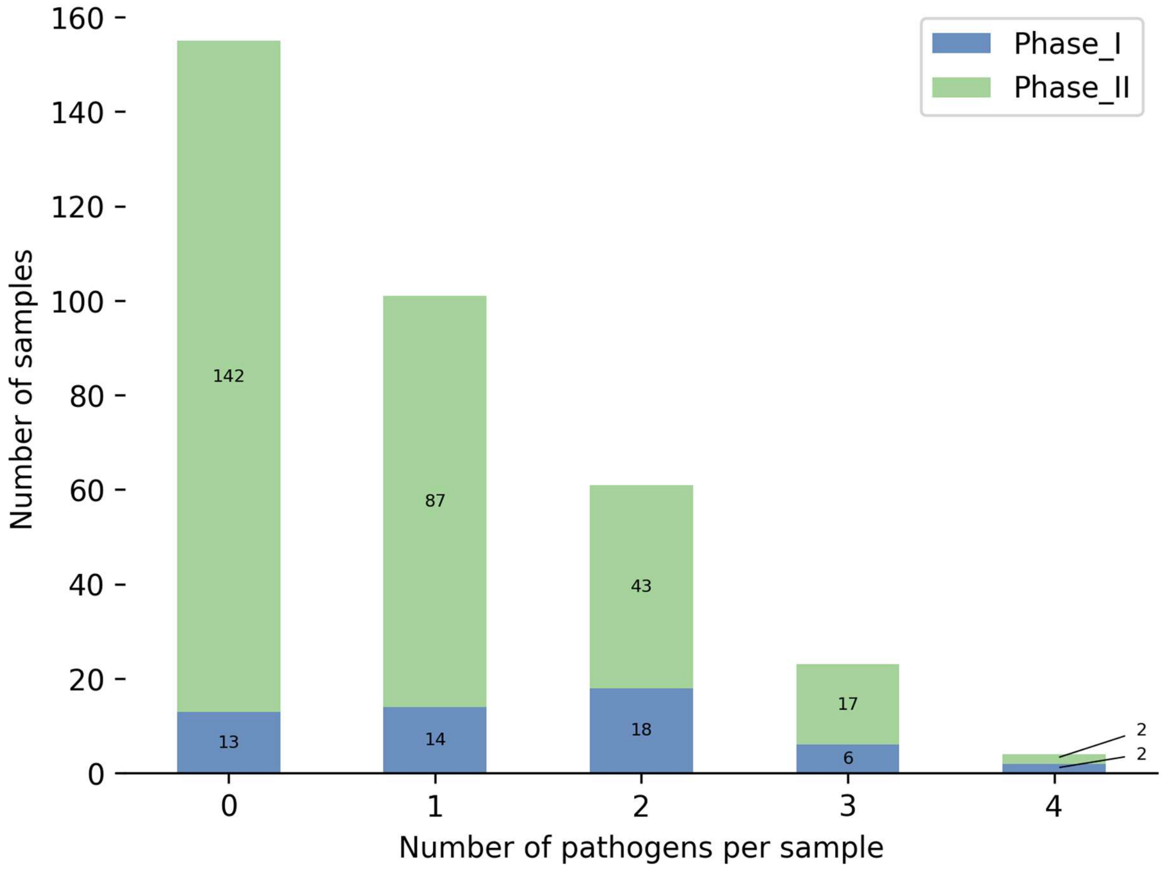


**Supplementary Figure 1: Summary of the number of pathogens identified among the clinical samples.** The histogram shows the number of pathogens identified in the phase II and phase I samples using routine microbiology with MALDI-TOF. A value of 0 represents the number of culture-negative samples (155); samples with one pathogen (101) were mono-microbial, while those with more than one pathogen were considered polymicrobial (88). Five samples from phase II were inconclusive due to sampling errors.
