## Supplementary Figure 2 for "Rapid clinical metagenomics enables early tailored therapy in complicated urinary tract infections and strengthens antimicrobial stewardship"

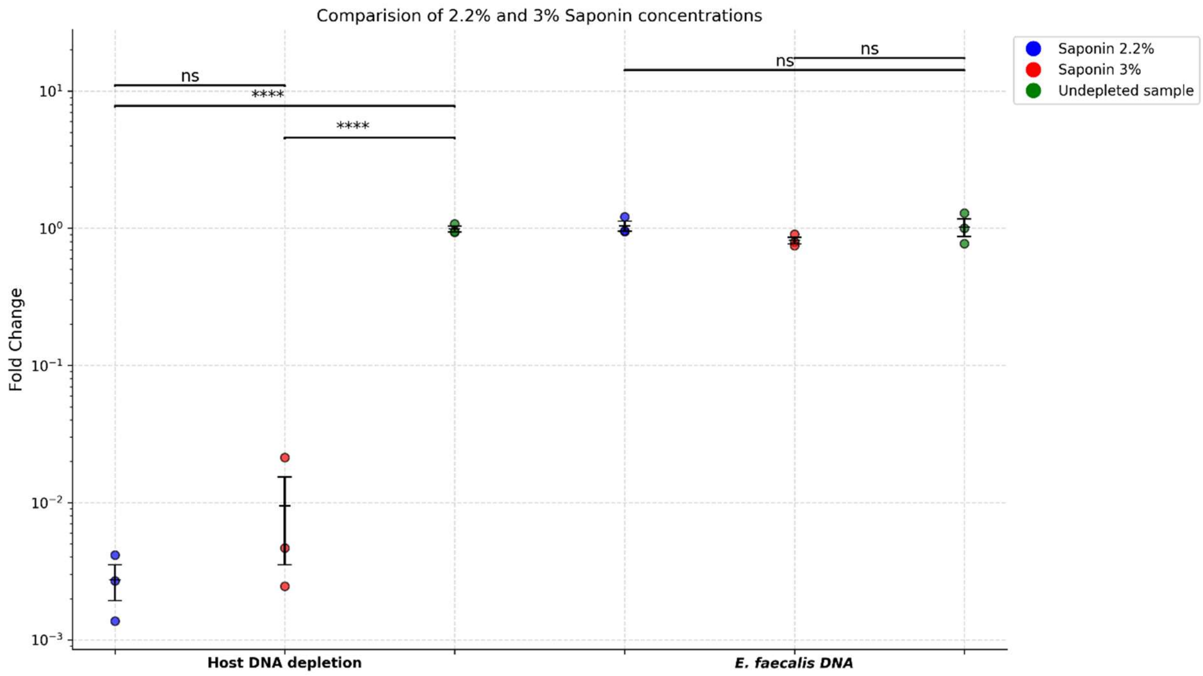


**Supplementary Figure 2: Figure comparing the effect of 2.2% and 3% saponin on the host depletion.** The samples were spiked with E. faecalis at $10^5$ CFU/mL along with the WBCs isolated from the whole blood at $1x10^4$ cells/mL as shown previously. No significant changes in the extent of host depletion were observed with changes in saponin concentration. All test values are mean ± SD, with n=3 biological replicates for each condition. ns > 0.05,*p≤0.05,**p≤0.01,***p≤0.001,****p≤0.0001.
