## Supplementary Figure 3 for "Rapid clinical metagenomics enables early tailored therapy in complicated urinary tract infections and strengthens antimicrobial stewardship"

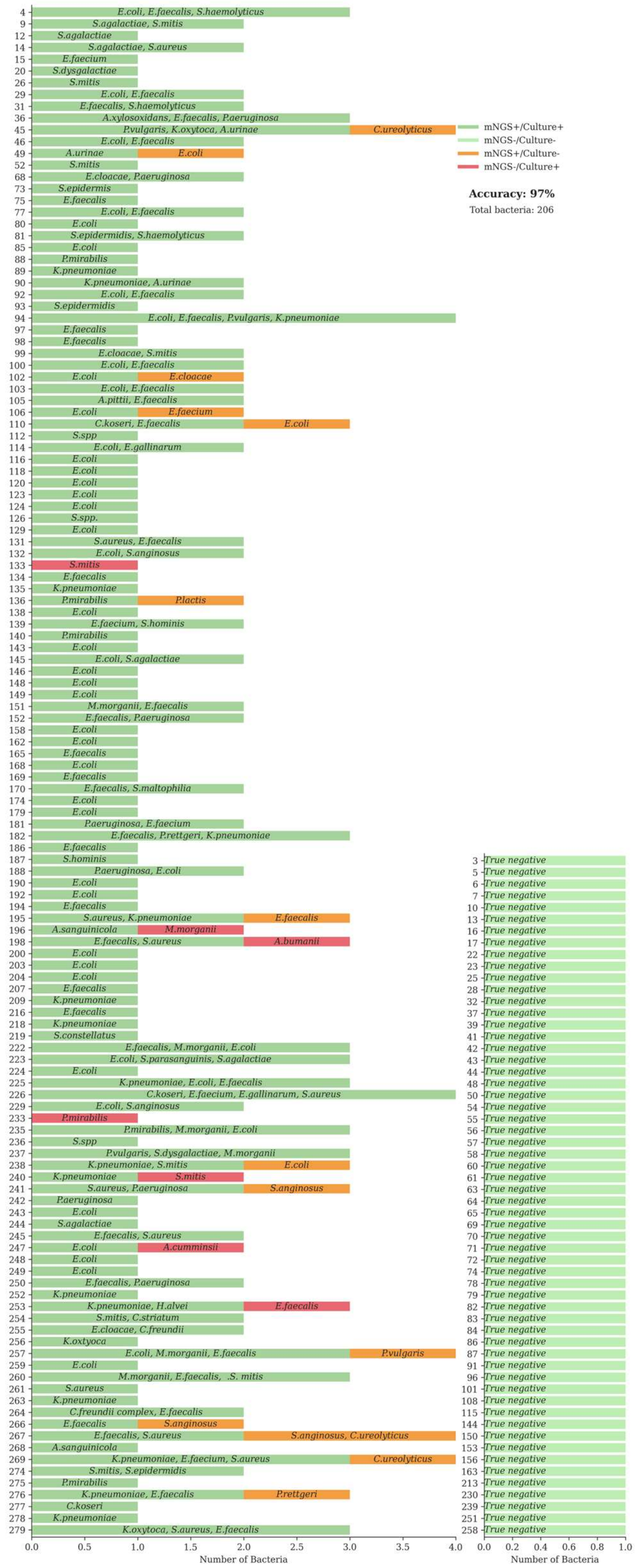


**Supplementary Figure 3: Pathogen identification results stratified by the samples.** The figure summarizes benchmarking results for pathogen identification from routine testing using MALDI-TOF and the URINN method. Dark green bars represent species identified by both methods (concordant). Light green bars represent samples that were negative by both methods (concordant). Orange bars represent species uniquely identified by the URINN method, while red bars indicate species identified through routine testing.
