## Supplementary Figure 4 for "Rapid clinical metagenomics enables early tailored therapy in complicated urinary tract infections and strengthens antimicrobial stewardship"

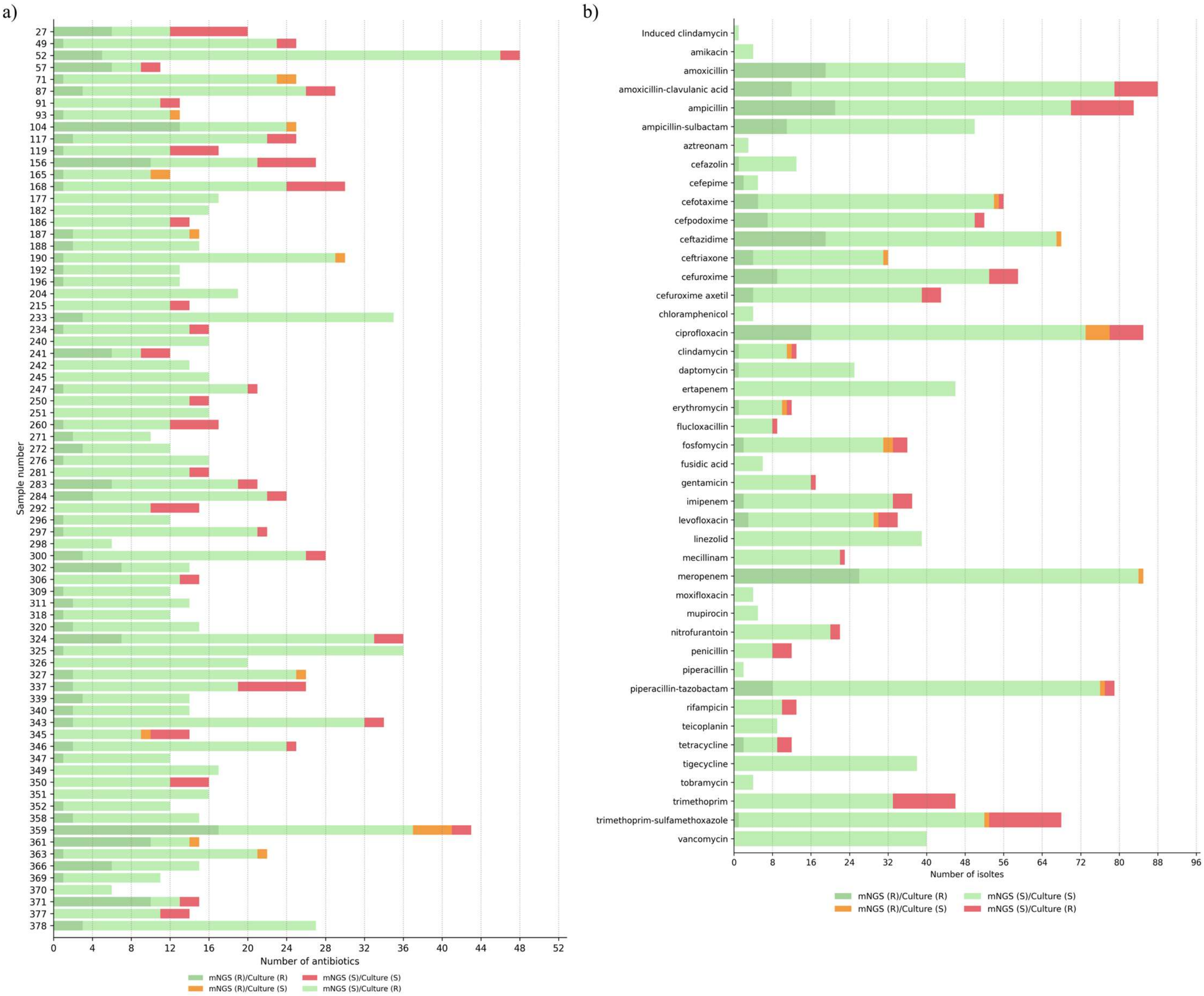


**Supplementary Figure 4: AST benchmarking concordance stratified by the sample and antibiotics**. The subfigure (a) represents the sample level concordance, while the subfigure (b) represents the antibiotic level concordance between the routine antimicrobial susceptibility testing performed using VITEK-2 and metagenomics (mNGS) resistance predictions obtained from the antibiotic resistance gene data. Green bars indicate concordance between observed phenotypes and detected ARGs (light green indicates true susceptibility and dark green indicates true resistance). Red bars highlight instances in which resistance was identified in routine AST but no corresponding resistance mechanism was detected by mNGS (false negatives). Orange bars signify detected ARGs without phenotypic resistance to the corresponding antibiotic. AMR predictions were made exclusively for antibiotics present in both the phenotypic and genotypic datasets.
