## Supplementary Figure 5 for "Rapid clinical metagenomics enables early tailored therapy in complicated urinary tract infections and strengthens antimicrobial stewardship"

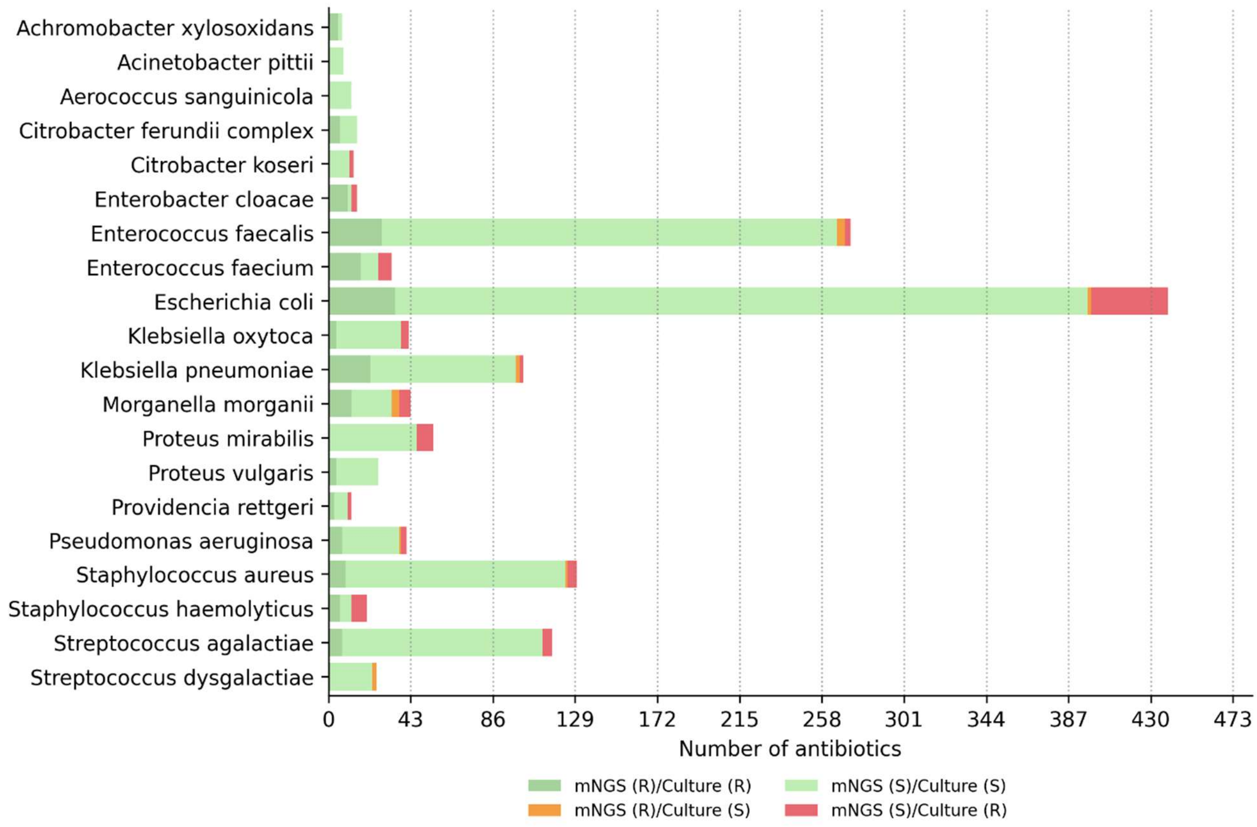


**Supplementary Figure 5: AST benchmarking concordance stratified on the pathogen level**. The figure shows the pathogen-level concordance between routine antimicrobial susceptibility testing performed using VITEK-2 and metagenomics (mNGS)- based resistance predictions derived from antibiotic resistance gene data. Green bars indicate concordance between observed phenotypes and detected ARGs (light green indicates true susceptibility and dark green indicates true resistance). Red bars highlight instances in which resistance was identified in routine AST, but no corresponding resistance mechanism was detected by mNGS (false negatives). Orange bars signify detected ARGs without phenotypic resistance to the corresponding antibiotic. AMR predictions were made exclusively for antibiotics present in both the phenotypic and genotypic datasets.
