## Supplementary Figure 6 for "Rapid clinical metagenomics enables early tailored therapy in complicated urinary tract infections and strengthens antimicrobial stewardship"

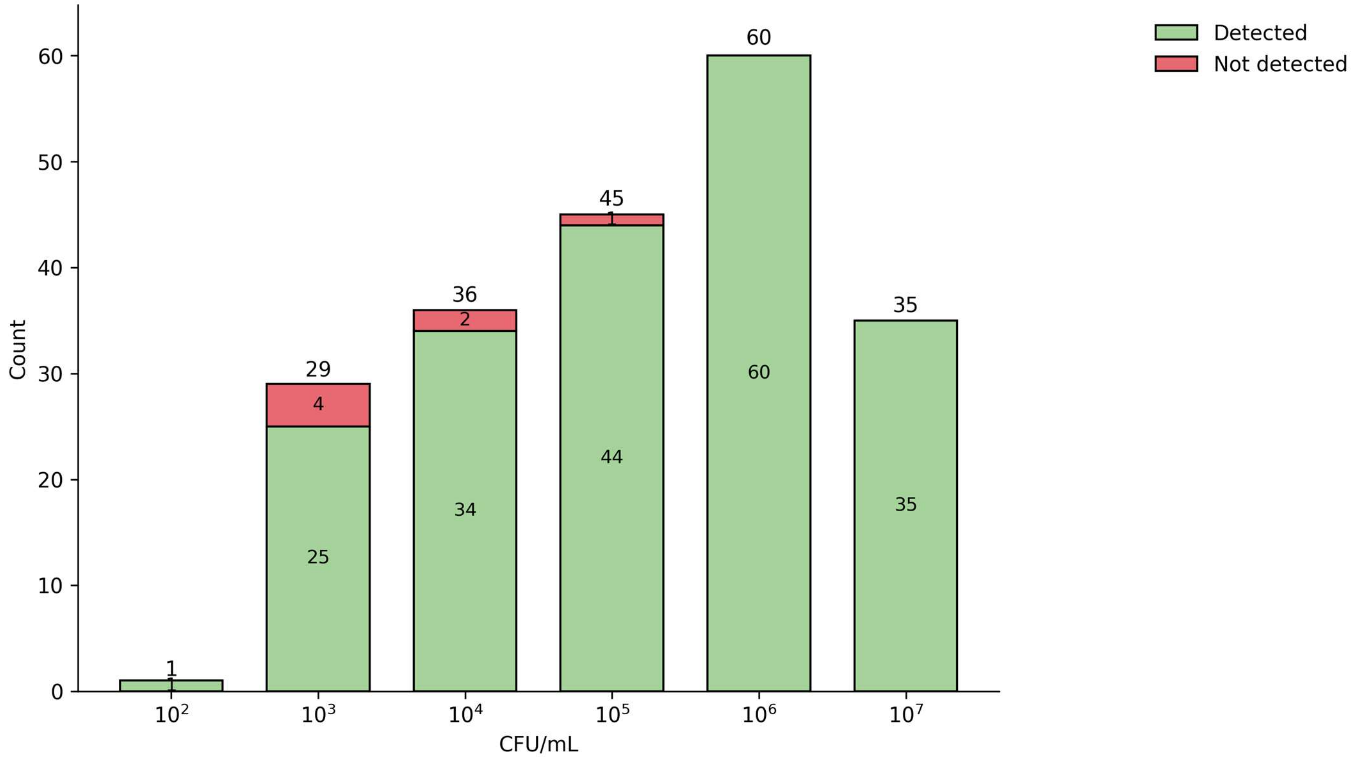


**Supplementary Figure 6: Stacked bar graph showing detected and missed pathogen species based on CFU/mL.** The figure presents a stacked bar graph illustrating detected and missed pathogen species across different CFU/mL concentrations. The figure conveys the number of pathogen species identified at each CFU/mL level within the phase II cohort. Green bars indicate detected species, while red bars represent missed pathogens. The numerical value above each bar denotes the total number of pathogens—both detected and missed—at that specific concentration.
