## Supplementary Figure 7 for "Rapid clinical metagenomics enables early tailored therapy in complicated urinary tract infections and strengthens antimicrobial stewardship"

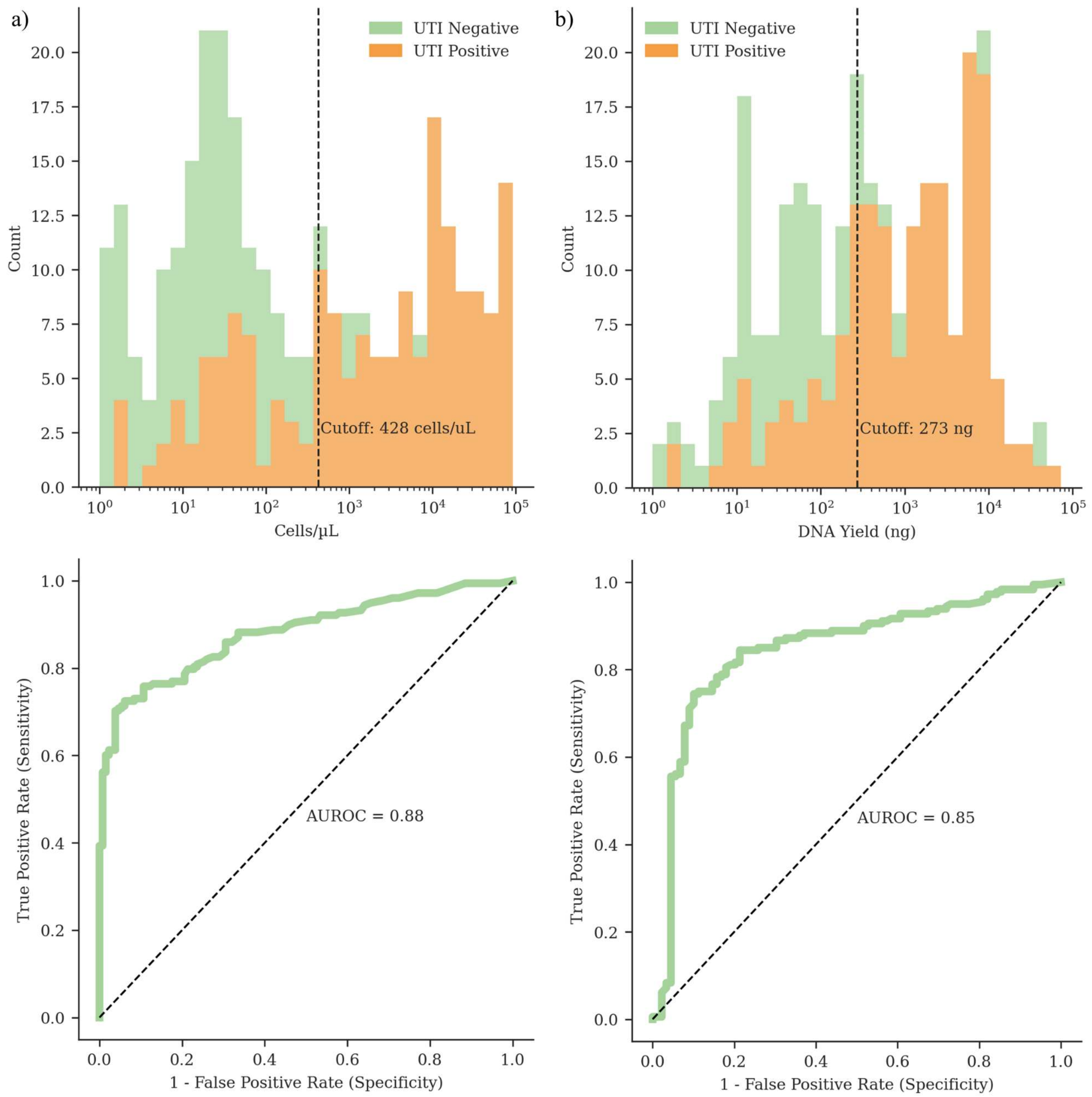


**Supplementary Figure 7: Bacterial cell and DNA yield receiver operating characteristic curve analysis.** Subfigure (a) displays the results for the number of bacterial cells/µL as identified through flow cytometry, and (b) presents the results for DNA yield. The analysis utilized available data from 309 samples for bacterial cell numbers and 270 samples for DNA yield. 35 samples were excluded from phase II due to a lack of flow cytometry data or inconclusive culture results, while 5 samples (1 culture positive and 4 culture negative) from Phase I were excluded due to the lack of flow cytometry data. For DNA analysis, 53 samples from phase I and 217 samples from phase II were used. Two samples from phase II were excluded because the DNA concentration was too low for quantification. For bacterial cell-based analysis, Phase II data were available for 288 samples (139 culture-positive and 114 culture-negative), and 53 samples (39 culture-positive and 9 culture-negative) were from Phase I. The histogram (top) illustrates the distribution of each variable for culture-positive and culture-negative samples, highlighting the optimal cutoff for distinguishing between the two groups. The ROC curve (bottom) is annotated with the area under the ROC curve (AUROC) for each variable.
