## Supplementary Figure 8 for "Rapid clinical metagenomics enables early tailored therapy in complicated urinary tract infections and strengthens antimicrobial stewardship"

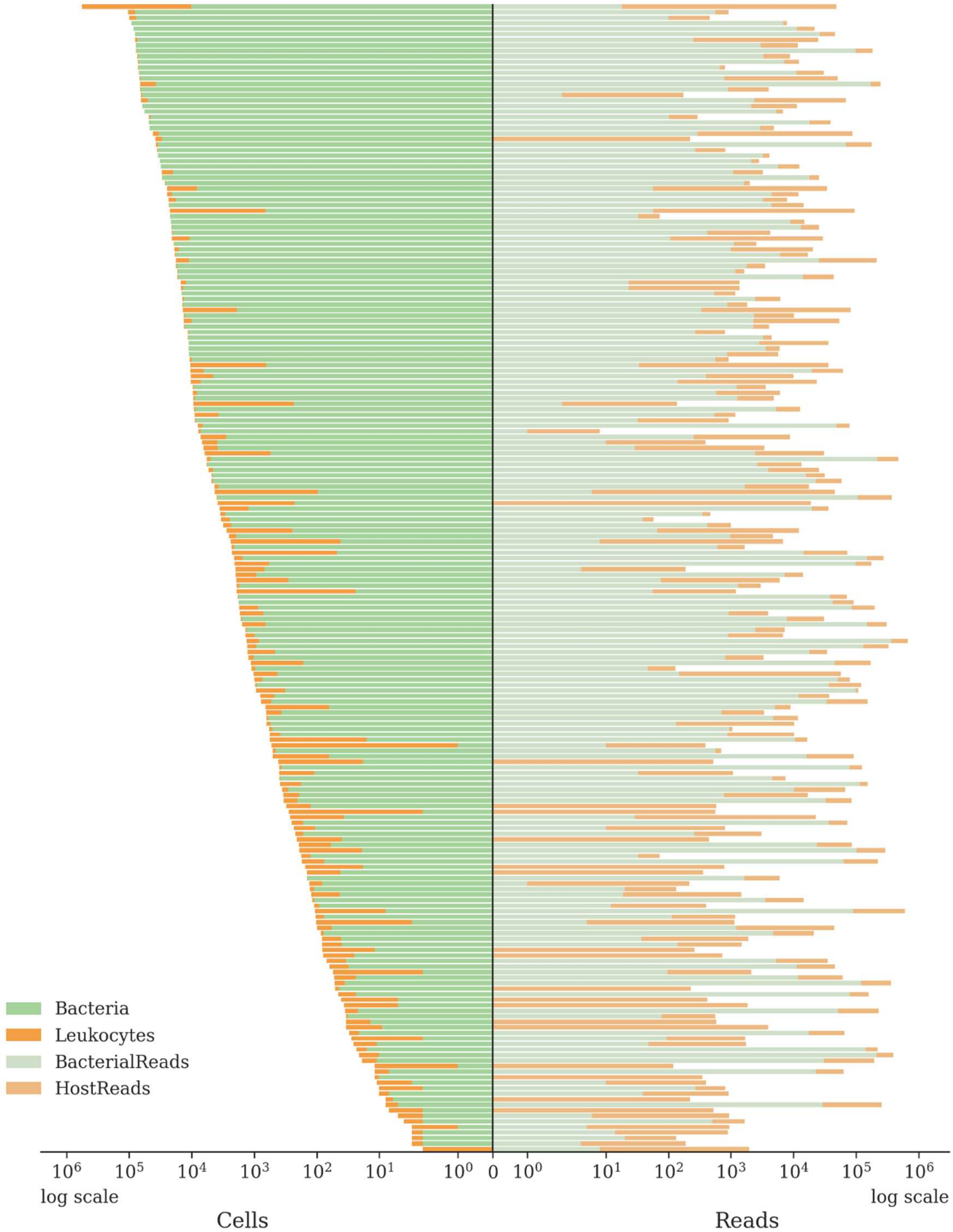


**Supplementary Figure 8: Correlation between flow cytometry and mNGS abundance.** The tornado charts illustrate the relationship between leukocytes (in orange) and bacterial cells (in green) based on flow cytometry data (left) and host-to-bacterial reads (right) obtained through metagenomic next-generation sequencing (mNGS). The data are visualized on a log scale. These plots enhance our understanding of the effectiveness of various methodologies across a range of clinical samples for the URINN method.
