## Supplementary Figure 9 for "Rapid clinical metagenomics enables early tailored therapy in complicated urinary tract infections and strengthens antimicrobial stewardship"

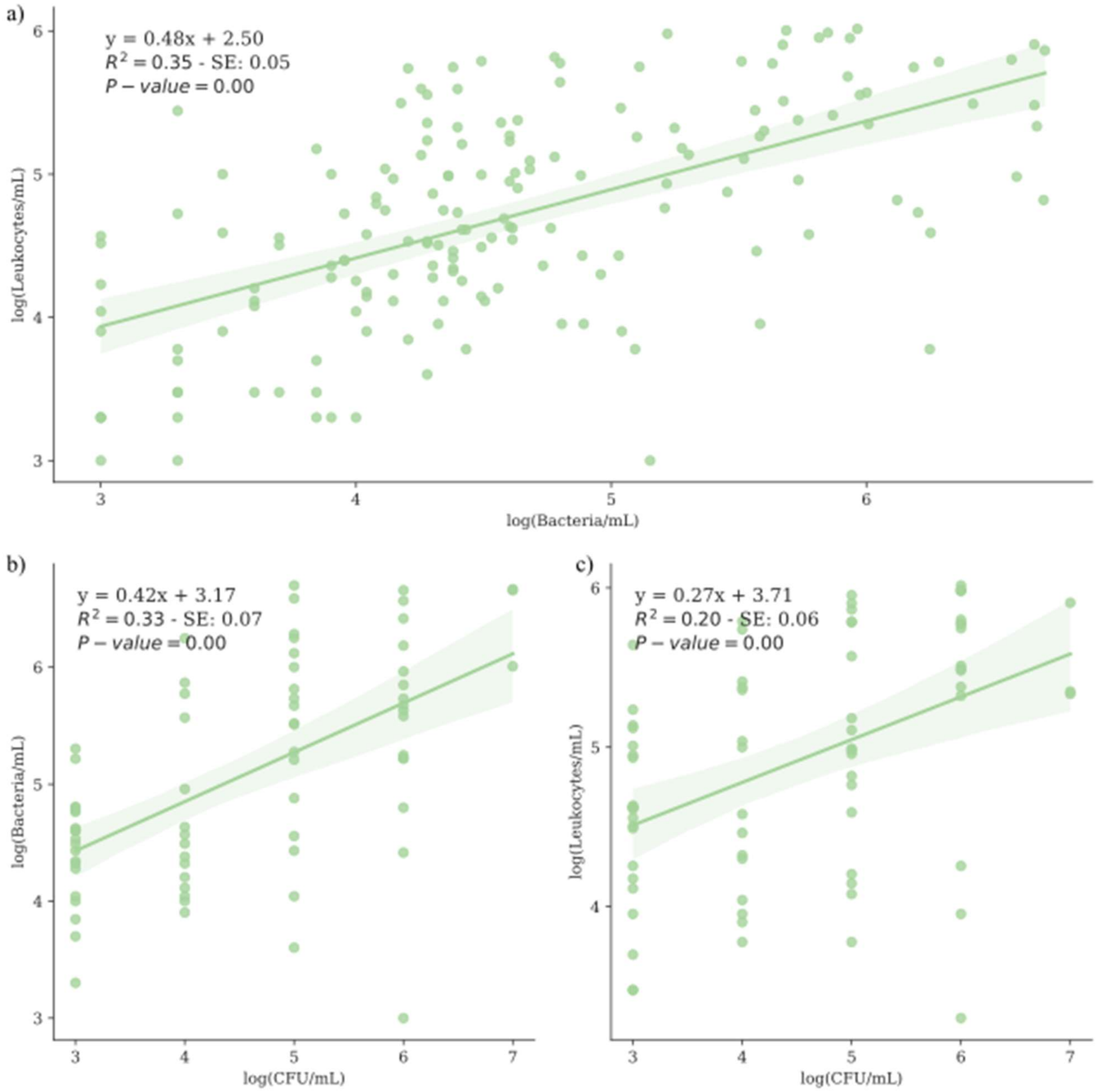


**Supplementary Figure 9: Linear regression of leukocyte and bacterial cell count and colony-forming units (CFU/mL).** Linear regression was performed after removing outliers using the interquartile range (IQR) method with lower bound (Q1 – 1.5 IQR) and upper bound (Q3 + 1.5 IQR). The plots were annotated with the linear model function (y), the coefficient of determination ($R^2$), the standard error of the model slope (SE), and the P-value obtained using the Wald Test with t-distribution of the test statistic. Individual samples are plotted as points in the scatter plot, and the regression line is plotted with its 95% confidence interval (shaded area) calculated using bootstrapping. The regression plots are shown with logarithmic axes. Subplot a) shows the linear regression of leukocyte to bacterial cell numbers obtained using flow cytometry. Regression of colony-forming units (CFU) to numbers obtained using flow cytometry is shown for bacterial cells b) and leukocytes c).
