## Supplementary Figure 10 for "Rapid clinical metagenomics enables early tailored therapy in complicated urinary tract infections and strengthens antimicrobial stewardship"

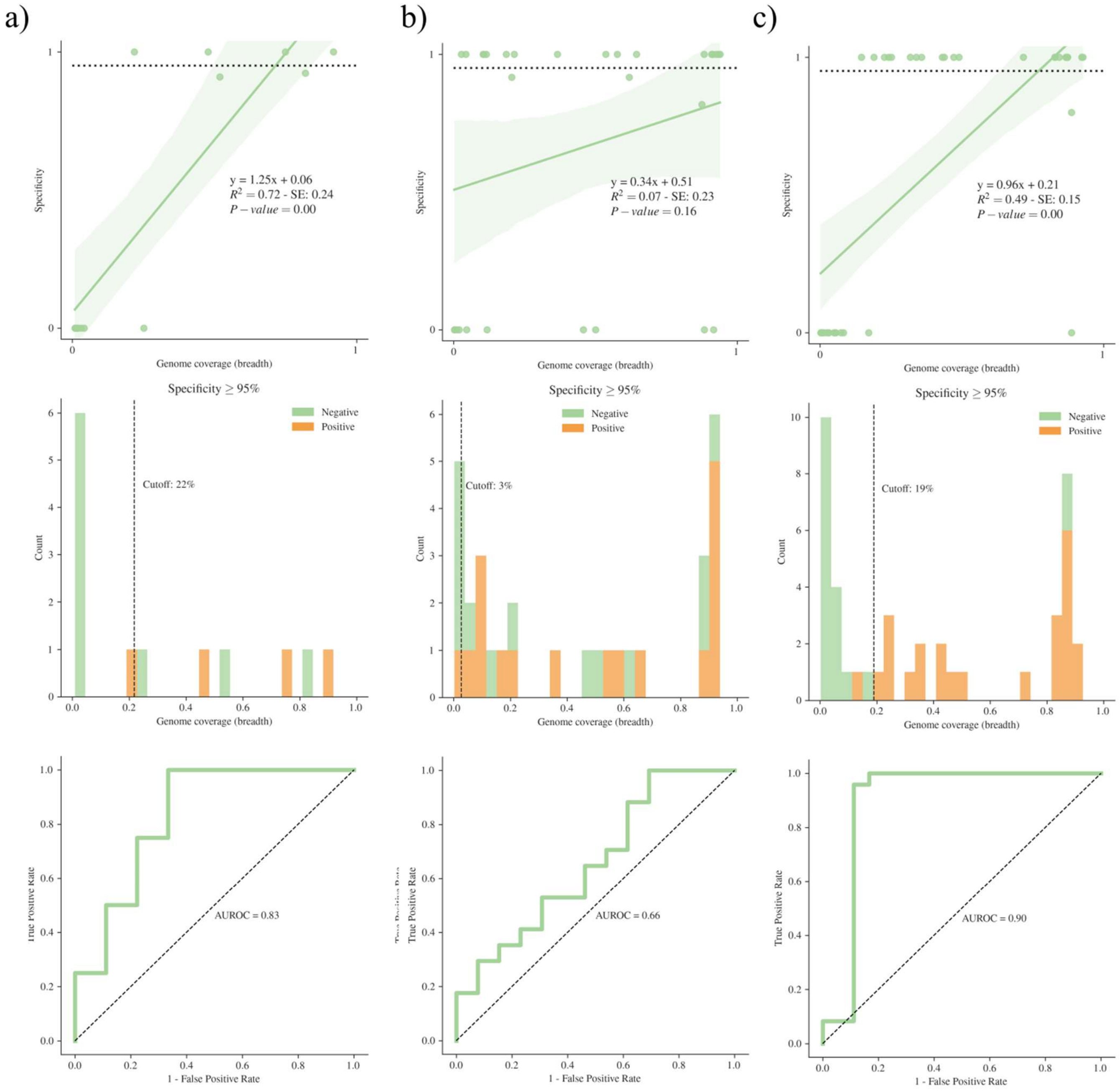


**Supplementary Figure 10: Correlation between AST prediction specificity and breadth of genome coverage.** Linear regression and AUROC analysis show the relationship between genome coverage and AST prediction specificity for a) *Klebsiella pneumoniae*, b) *Enterococcus faecalis*, and c) *Escherichia coli*. Linear regression plots (top) show the linear correlation between breadth of genome coverage and AST prediction sensitivity, where each data point represents an individual pathogen. The regression function (y), the coefficient of determination ($R^2$), the standard error of the model slope (SE), and the P-value are reported. P-values were computed using the Wald Test with the t-distribution. The histogram (center) shows the distribution of pathogens with AST prediction specificity above (Positive) or below (Negative) 95%, and indicates the optimal cutoff determined by minimizing the Youden's index. The AUROC curve (bottom) is annotated with the area under the ROC curve.
