## Supplementary Figure 11 for "Rapid clinical metagenomics enables early tailored therapy in complicated urinary tract infections and strengthens antimicrobial stewardship"

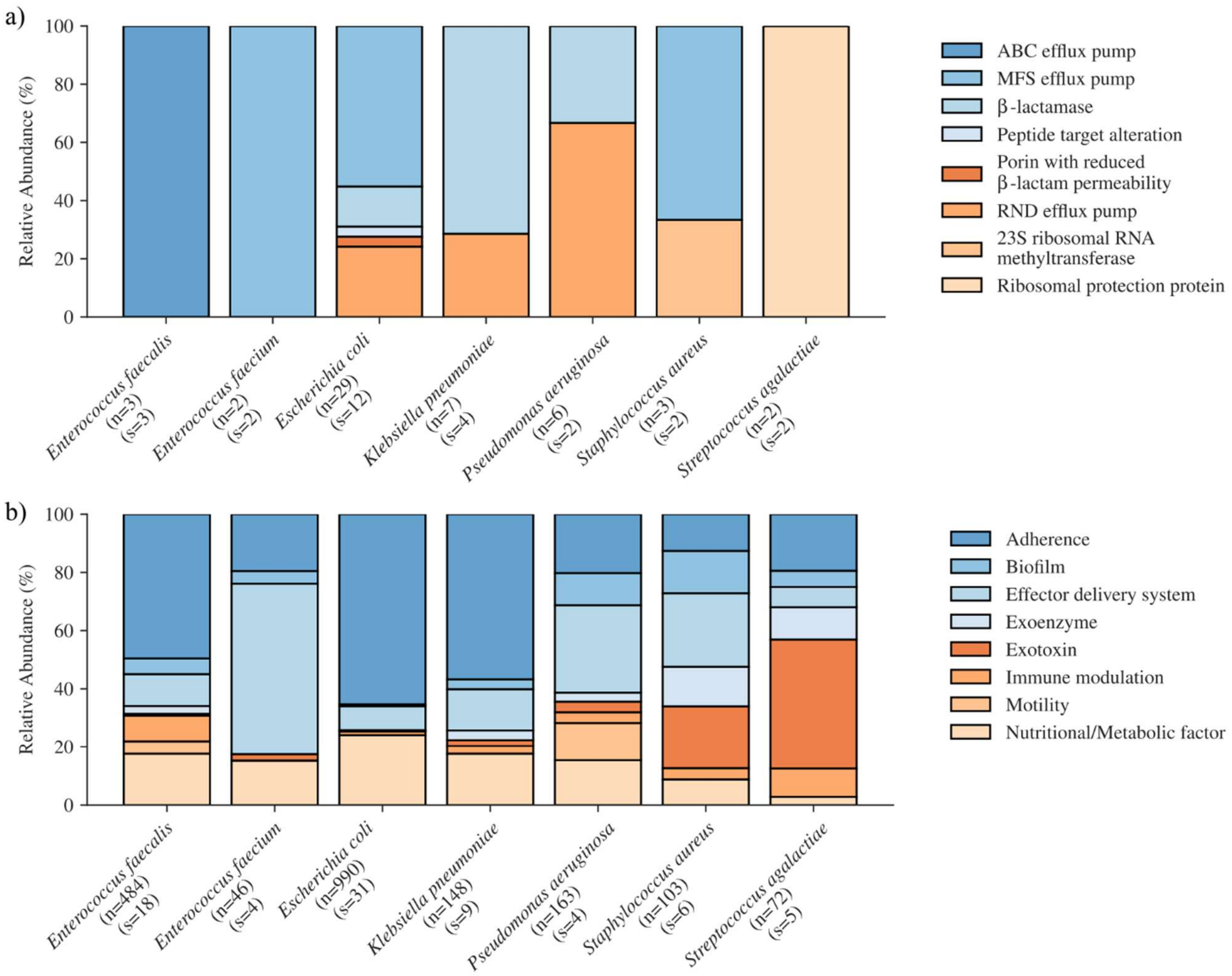


**Supplementary Figure 11: Comparison of resistance and virulence mechanisms per species.** The relative abundance of a) resistance mechanisms and b) virulence mechanisms identified through mNGS is shown grouped by species. The species names are annotated with the total number of identified genes (n) and the number of unique samples (s) from which the data were derived. Abbreviations: ATP-binding cassette (ABC), major facilitator superfamily (MFS), resistance-nodulation-cell division (RND).
