## Supplementary Table 1 for "Rapid clinical metagenomics enables early tailored therapy in complicated urinary tract infections and strengthens antimicrobial stewardship"

| | | Total number of samples | Samples with DNA extracted | Number of samples sequenced |
|---|---|---|---|---|
| 349 patient samples | Phase I | 53 (40/13) | 53 (40/13) | 53 (40/13) |
| | Phase II | 296 (149/142/5) | 219 (141/77/1) | 189 (130/58/1) |

**Supplementary Table 1:** Sample distribution table summarizes the distribution of samples at different stages of processing, based on two testing phases, I and II. The figures in brackets represent the number of culture-positive samples (highlighted in green), culture-negative samples (highlighted in red), and samples excluded from analysis due to sampling errors, such as sample spillage and inconclusive data from routine results (highlighted in blue).
